# Polygenic prediction across populations is influenced by ancestry, genetic architecture, and methodology

**DOI:** 10.1101/2022.12.29.522270

**Authors:** Ying Wang, Masahiro Kanai, Taotao Tan, Mireille Kamariza, Kristin Tsuo, Kai Yuan, Wei Zhou, Yukinori Okada, the BioBank Japan Project, Hailiang Huang, Patrick Turley, Elizabeth G. Atkinson, Alicia R. Martin

**Author notes:** Lead contact: Ying Wang.

## Abstract

Polygenic risk scores (PRS) developed from multi-ancestry genome-wide association studies (GWAS), PRS_multi_, hold promise for improving PRS accuracy and generalizability across populations. To establish best practices for leveraging the increasing diversity of genomic studies, we investigated how various factors affect the performance of PRS_multi_ compared to PRS constructed from single-ancestry GWAS (PRS_single_). Through extensive simulations and empirical analyses, we showed that PRS_multi_ overall outperformed PRS_single_ in understudied populations, except when the understudied population represented a small proportion of the multi-ancestry GWAS. Notably, for traits with large-effect ancestry-enriched variants, such as mean corpuscular volume, using substantially fewer samples from Biobank Japan achieved comparable accuracies to a much larger European cohort. Furthermore, integrating PRS based on local ancestry-informed GWAS and large-scale European-based PRS improved predictive performance in understudied African populations, especially for less polygenic traits with large ancestry-enriched variants. Our work highlights the importance of diversifying genomic studies to achieve equitable PRS performance across ancestral populations and provides guidance for developing PRS from multiple studies.

## Introduction

Polygenic risk scores (PRS) have emerged as useful tools for estimating the cumulative genetic susceptibility to complex traits and diseases. PRS are typically calculated by weighting the number of risk alleles based on their associations in genome-wide association studies (GWAS). PRS have shown promising potential in predicting some traits and disease risks, comparable to monogenic variants and traditional clinical risk factors^1–5^. Achieving the most accurate and generalizable PRS requires access to large-scale and diverse GWAS, especially with representation that matches the specific target population. However, the current landsca pe of GWAS predominantly focuses on European (EUR) ancestry populations, which have considerably larger sample sizes compared to other populations. Although ongoing efforts are underway to rectify these gaps, achieving global representativeness is a challenging goal. Encouragingly, studies have shown that using GWAS data with even a small proportion of non-European ancestry individuals has the potential to improve the predictive accuracy of PRS in underrepresented populations^6–8^. This finding could largely be attributed to the substantial contribution of common variants to the heritable variation of complex traits and diseases, and that causal variants are largely shared across ancestries^9–12^. With the ever-increasing availability and scalability of genomic data from underrepresented and ancestrally diverse populations, we are especially interested in leveraging this greater diversity to improve PRS generalizability.

In particular, recently admixed populations, consisting of chromosomal segments of mosaic ancestries, are systematically excluded in many existing genomic studies due to their underrepresentation and complicated population structure^13–15^. However, these populations present unique opportunities to develop more generalizable PRS as their genetic effects can be estimated in more consistent environments, which helps reduce confounding factors compared to estimates across different ancestry groups in different populations^16^. Furthermore, the comprehensive characterization of phenotypes is often insufficient or inconsistently performed in different populations. However, in the recently admixed populations, there is a greater potential for consistency and comparability in phenotype measurements, as the genetic factors contributing to phenotypic differences between the source populations can be decoupled in the recently admixed populations^16, 17^. The advancement of methodologies such as local ancestry inference and association testing has further enabled ancestry-specific GWAS in admixed populations^18–20^, allowing for the construction of PRS that leverage genetic information captured by local ancestry inference. With the ongoing accumulation of data from recently admixed populations, particularly through initiatives like the *All of Us* Research Program^21^, expanded resources will provide unparalleled opportunities to explore the performance of PRS derived from local ancestry-informed summary statistics within historically underrepresented populations. Furthermore, such data will facilitate their integration with PRS derived from predominantly EUR-based cohorts.

Recently developed statistical methodologies leverage the increasing diversity of GWAS data to improve PRS portability^8, 22, 23^. However, the effect of genetic architecture, ancestry composition of GWAS discovery cohorts, and PRS construction methodologies on cross-ancestry predictive accuracy remains largely unclear. For example, a recent study found no increase in accuracy when meta-analyzing GWAS from a relatively small Ugandan cohort with larger EUR data^24^. Furthermore, theoretical frameworks for approximating expected PRS accuracy from multi-ancestry GWAS are lacking. Current theoretical calculations for PRS accuracy rely on the assumption of homogeneity within the ancestral discovery samples^25, 26^, ignoring factors that are likely to play a role with multi-ancestry cohorts. Such factors may include differences in linkage disequilibrium (LD), minor allele frequency (MAF), heritability, sample sizes, and genetic correlation across different ancestries.

To provide insights into those issues, we explored the impact of ancestry compositions in discovery GWAS on predictive accuracy of PRS constructed using different methodologies. This exploration involved large-scale population genetic simulations as well as the utilization of real genomic data from the BioBank Japan (BBJ)^27^ and UK Biobank (UKBB)^6^ across traits exhibiting distinct genetic architectures (**Figure 1)**. In what follows, we used **single-ancestry GWAS** to denote studies conducted exclusively within a single ancestry group (defined using genetic data), while **multi-ancestry GWAS** refers to studies encompassing two or more distinct ancestries. In our analyses, we performed meta-analyses of GWAS conducted in European ancestry populations (**EUR GWAS)** and GWAS conducted in other minority populations (**Minor GWAS**) by varying the ratios of sample sizes to mimic multi-ancestry GWAS with varying ancestry compositions. Specifically, we focused on East-Asian (EAS) and African (AFR) minority populations. By comparing the performance of PRS derived from single-ancestry GWAS (referred to as **PRS_single_**) and multi-ancestry GWAS (referred to as **PRS_multi_**) through simulations and real data, we consistently observed that PRS_multi_ overall exhibited superior performance in comparison to PRS_single_ (primarily PRS derived from large-scale EUR GWAS, referred to as **PRS_EUR_GWAS_**). As admixed populations remain understudied despite disproportionately yielding novel genetic findings^28^, we further conducted local ancestry inference to explore whether, how, and to what extent PRS performance could be improved using GWAS discovery data from AFR-EUR admixed individuals. While optimal PRS methods are trait- and context-specific, this study comprehensively evaluates PRS accuracy across a wide range of scenarios, facilitating a set of best practices that ultimately reduces the number of analyses necessary to optimize PRS for specific applications.

**Figure 1.**
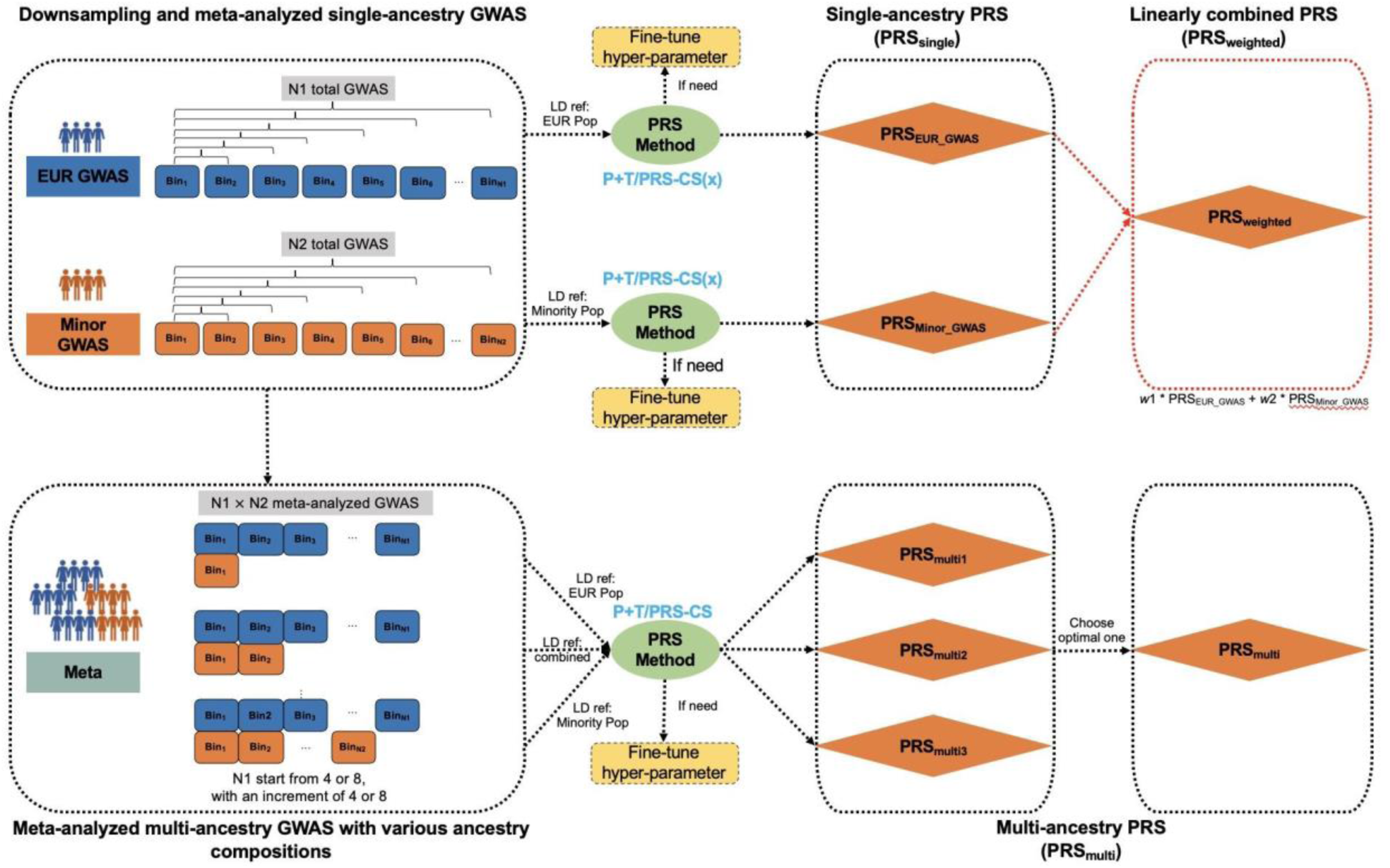
Study design in both simulations and empirical analyses. 1) In the context of single-ancestry GWAS, we randomly split individuals in European (EUR) and other minority populations, including East-Asian and African populations, into equally sized bins. Simulations involved a total of 52 bins per population, each containing 10,000 individuals. For empirical analysis, bin number was dependent on the sample size of respective phenotype in that population (**Table S3**), with 5,000 individuals per bin. GWAS was conducted within each bin for each individual population, followed by meta-analysis of GWAS from various numbers of bins within each population. To construct PRS derived from single-ancestry GWAS (**PRS_single_**) in the target population, we applied P+T for both simulations and empirical analyses, utilizing PRS-CS for the latter. Subsequently, we combined PRS_single_ developed from EUR GWAS (**PRS_EUR_GWAS_**) and other minority population-based GWAS (**PRS_Minor_GWAS_**) through a linear weighted strategy (denoted as **PRS_weighted_**, highlighted in red box) for empirical analyses. Note that PRS_weighted_ was also developed using PRS-CSx, which utilizes GWAS summary statistics from multiple populations. 2) For meta-analyzed multi-ancestry GWAS (referred to as **Meta),** we ran meta-analyses on EUR GWAS and Minor GWAS with varying ancestry compositions. In simulations, we incrementally included 4 bins from EUR GWAS for the meta-analysis, while in empirical analyses, we increased the number to 8 bins. Simultaneously, we varied the number of bins in Minor GWAS from 1 to the total number. Following the meta-analysis, we constructed PRS based on Meta (referred to as **PRS_multi_**), using the P+T method for simulations, and employing both P+T and PRS-CS for empirical analyses.

## Results

### Evaluating the effects of imbalanced sample sizes across ancestries on PRS accuracy through simulations

We simulated genotypes using HapGen2 and phenotypes according to six different scenarios with varying trait heritability (*h*^2^ = 0.03, 0.05) and number of causal variants *(*M*_*c*_* = 100, 500, 1000), such that the polygenicity ranged from ∼0.1% to ∼1%. We assumed that the causal variants and their effect sizes are shared across ancestries (i.e., cross-ancestry genetic correlation, *r_g_*, is 1) in our initial simulations. For single-ancestry GWAS, we first conducted GWAS within each bin and then meta-analyzed GWAS across different numbers of bins (1-52 per ancestry). Each bin represented 10,000 individuals randomly sampled from the corresponding ancestry. For multi-ancestry GWAS, we meta-analyzed GWAS from EUR and minor populations (EAS or AFR) to evaluate the impact of ancestry composition. We used varying numbers of bins from the EUR GWAS (ranging from 4 to 52 with 4 increments) and varied the contribution from minority populations (1-52 bins) from EAS or AFR GWAS. We constructed PRS using the classic pruning and thresholding (P+T) method with varying *p*-value thresholds. This approach follows a greedy heuristic algorithm wherein variants are sorted based on their *p*-values. The algorithm iteratively descends in significance while retaining only those variants that do not exceed a predetermined LD threshold with previously retained variants. We assessed the accuracy, measured by prediction *R*^2^, using the optimal threshold through fine-tuning in the validation cohort. Detailed information about the simulation setup is shown in **Figure 1** and **STAR Methods**.

### PRS predictive accuracy improved with more individuals from target populations included in the multi-ancestry GWAS but varied with genetic architecture

When developing PRS using single-ancestry GWAS, we found that using ancestry-matched GWAS generally outperformed using GWAS from other discovery populations (**Figure S1**). Compared to using EUR GWAS, the benefit of using ancestry-matched GWAS was more evident for traits with more polygenic genetic architectures and larger GWAS sample sizes. To further evaluate the impact of ancestry composition, we compared the accuracy of PRS_multi_ and PRS_single_. We constructed PRS_multi_ using an LD reference panel consisting of individuals proportional to the ancestry composition of the discovery GWAS. This reference panel yielded approximately optimal accuracy among three different reference panels utilized in our study (**Figure S2 and Supplementary Note 1, 2**).

Relative to the accuracy of PRS_EUR_GWAS_, we observed significant improvements in the understudied target population by including more individuals from the target ancestry in multi-ancestry GWAS. Across all simulations, a statistically significant median improvement of 0.008 in *R*^2^ was observed (one-sided Wilcoxon signed-rank test, *p*-value < 2.2e-16, **Table S1**). This trend was more apparent in more polygenic traits. As shown in **Figure 2**, we compared accuracy between PRS_multi_ and PRS_EUR_GWAS_ derived from 320,000 EUR individuals. For traits with *h*^2^ of 0.05, the median improvements in *R*^2^ of PRS_multi_ was 0.006, 0.014 and 0.013 with *M_c_* of 100, 500, and 1000, respectively, in the EAS target population. Similarly, corresponding *R*^2^ improvements of 0.009, 0.010 and 0.014 were shown in AFR (**Figure S3**). However, we did not consistently observe such accuracy gains for the majority EUR population, or in scenarios where the other understudied ancestry was not included in the multi-ancestry discovery GWAS. In our simulations but unlike in most GWAS, populations typically understudied in current genomic studies can be the majority in the discovery GWAS. Nevertheless, we still observed significant PRS accuracy improvements, of median improvements in *R*^2^ 0.007 across simulations when the proportion of understudied populations in the discovery GWAS was less than 50% (one-sided Wilcoxon signed-rank test, *p*-value < 2.2e-16). We expected to observe similar relative *R*^2^ improvements, which measured the PRS generalizability, in the target populations using PRS_multi_ compared to using PRS_EUR_GWAS_ with the same number of bins from EUR populations (**Supplementary Note 3**).

**Figure 2.**
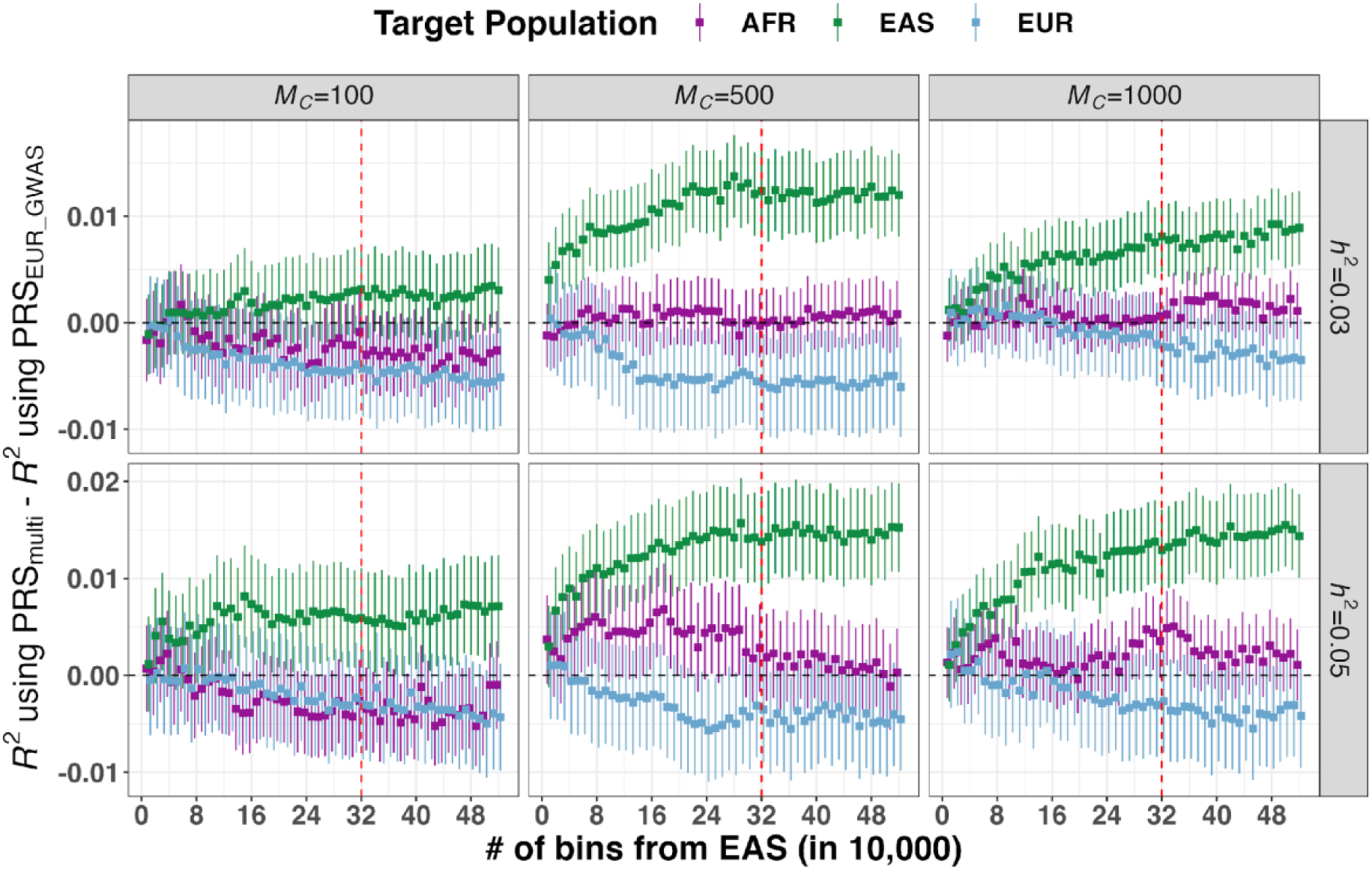
Improvement of PRS accuracy through meta-analyzed multi-ancestry GWAS compared to large-scale European GWAS across 6 simulated genetic architectures. The multi-ancestry GWAS included populations of European (EUR) and East-Asian (EAS) ancestry, with the EAS sample size varying as indicated on the x-axis. For illustrative purposes, we present the results using 32 EUR bins, each consisting of 10,000 individuals, which were included in both EUR GWAS and multi-ancestry GWAS. PRS was separately evaluated in African (AFR), EAS and EUR populations. Full results are shown in **Table S1**. *M_c_* indicates the number of causal variants and *ℎ*^2^ refers to SNP-based heritability. In each panel, the red vertical dashed line indicates the point where an equal number of bins from EUR and EAS populations were included in the multi-ancestry GWAS. The error bars represent the standard errors of predictive accuracy differences between PRS derived from multi-ancestry GWAS (PRS_multi_) and PRS derived from EUR GWAS (PRS_EUR_GWAS_).

Compared with using PRS_EUR_GWAS_, we found that PRS_multi_ derived from GWAS with much smaller sample sizes could achieve comparable or better predictive accuracy (**Table S1**). For example, in the scenario with *M_c_* of 1000 and *h*^2^ of 0.03, the meta-analysis of 16 EUR and 2 AFR bins achieved a comparable accuracy of 0.008 to that of using 32 EUR bins in the AFR population. Overall, adding fewer individuals from the target populations saturated accuracy improvements for less polygenic traits faster than more polygenic traits. Similarly, larger sample sizes from AFR populations were required to achieve comparable PRS accuracy to EAS populations especially for more polygenic traits, likely due to the larger effective population size in AFR populations and larger genetic divergence between EUR and AFR populations. As shown in **Figure S3**, when *h*^2^ was 0.03, the accuracy improvement of PRS_multi_ in the AFR population plateaued to ∼0.005 with 11 and 20 AFR bins for *M_c_* of 100 and 500, respectively, but continued to increase with more AFR bins for *M_c_* of 1000. Similarly, when *h*^2^ was 0.03, including 2 and 12 EAS bins in PRS_multi_ yielded an accuracy improvement of >0.005 in EAS for *M_c_* of 100 and 500, respectively (**Figure 2**). In comparison to PRS derived from Minor GWAS alone (PRS_Minor_GWAS_), we found that the accuracy improvement of PRS_multi_ gradually diminished as the sample size of Minor GWAS increased (**Figure S4 and Table S1**). We showed that for more polygenic traits, PRS_multi_ achieved little to no improvement when the understudied target populations accounted for more than half of the sample size in multi-ancestry GWAS (**Supplementary Note 4)**.

Because genetic correlation estimates between populations can be significantly less than 1, we also modified our simulations by varying the *r_g_* to be 0.6 and 0.8. We investigated two simulation scenarios that represent the extremes in per-variant variance explained: the least polygenic scenario 1 with *M_c_* = 100 and *h*^2^ =0.05, and the most polygenic scenario 2 with *M_c_* = 1000 and *h*^2^ =0.03 (**STAR Method**). Consistent with our previous findings, PRS_multi_ exhibited improved predictive accuracy in the target population when a greater number of individuals from the same ancestry were included, as compared to relying solely on large-scale EUR GWAS (**Figure S5-A, B)**. This improvement was more pronounced for scenario 2. Moreover, we needed a larger number of individuals from the target ancestry to saturate accuracy improvements in scenario 1 when *r_g_* was moderately reduced. Furthermore, as the sample sizes of the Minor GWAS increased and the values of *r_g_* decreased, the advantage of utilizing PRS_multi_ over PRS_Minor_GWAS_ diminished and eventually vanished (**Figure S5-C, D)**. Details are shown in **Table S2** and S**upplementary Note 5.**

### Empirical analysis of PRS accuracy within and across ancestries using 17 quantitative phenotypes

#### Genetic architecture of 17 studied phenotypes

To understand how trait genetic architecture influences predictive accuracy of PRS across ancestries, we conducted a comprehensive analysis involving 17 phenotypes in the UKBB and BBJ. Specifically, we estimated key parameters influencing different aspects of genetic architecture, including SNP-based heritability, polygenicity (the proportion of SNPs with nonzero effects) and a coefficient of negative selection (*S*, measuring the relationship between MAF and estimated effect sizes). To obtain these estimates, we employed a Bayesian method called summary-data-based BayesS (SBayesS), which leverages GWAS summary statistics as input data^29^.

The phenotypes included in this study varied widely in genetic architecture across these estimated parameters (**Figure 3**, **Table S3 and Table S4**). The polygenicity estimates spanned a broad range, from low values (0.001-0.005) for traits like mean corpuscular hemoglobin concentration (MCHC), basophil count (basophil), mean corpuscular hemoglobin (MCH), and mean corpuscular volume (MCV), to higher values (0.012-0.047) for traits such as height and body mass index (BMI). SNP-based heritability estimates similarly ranged from <0.1 for basophil and MCHC to 0.54 and 0.33 for height using UKBB and BBJ, respectively, regardless of polygenicity. The median *S* parameters were -0.63 and -0.47 using UKBB and BBJ, respectively. While the negative *S* values indicate negative selection (i.e., rarer variants have larger effects), it remains unclear to what degree population stratification could confound such estimates^30, 31^. We found that the polygenicity estimates using UKBB were mostly higher than those using BBJ, which could be due to the higher statistical power with larger sample sizes in the UKBB resulting in the detection of more variants with small effects. Similarly, we observed significantly higher SNP-based heritability in the UKBB compared to BBJ with the exception of MCHC and basophil, indicating possible phenotype heterogeneity between the two cohorts. These results are expected from the biobank designs, as BBJ is a hospital-based cohort with participants recruited with certain diseases, whereas UKBB is a population-based cohort with overall healthier participants and thus a wider range of natural variation in complete blood counts. This finding is also consistent with the previous study using estimates from LD score regression (LDSC) and stratified-LDSC^32^. Moreover, as described previously^32^, the estimated cross-ancestry genetic correlations between UKBB and BBJ for those traits were not statistically different from 1 (*p*-value > 0.05/17) except for a few including basophil (*r_g_* = 0.5945, SE = 0.1221), height (*r_g_* = 0.6932, SE = 0.0172), BMI, (*r_g_* = 0.7474, SE = 0.0230), diastolic blood pressure (DBP, *r_g_* = 0.8354, SE = 0.0509), and systolic blood pressure (SBP, *r_g_* = 0.8469, SE = 0.0430).

**Figure 3:**
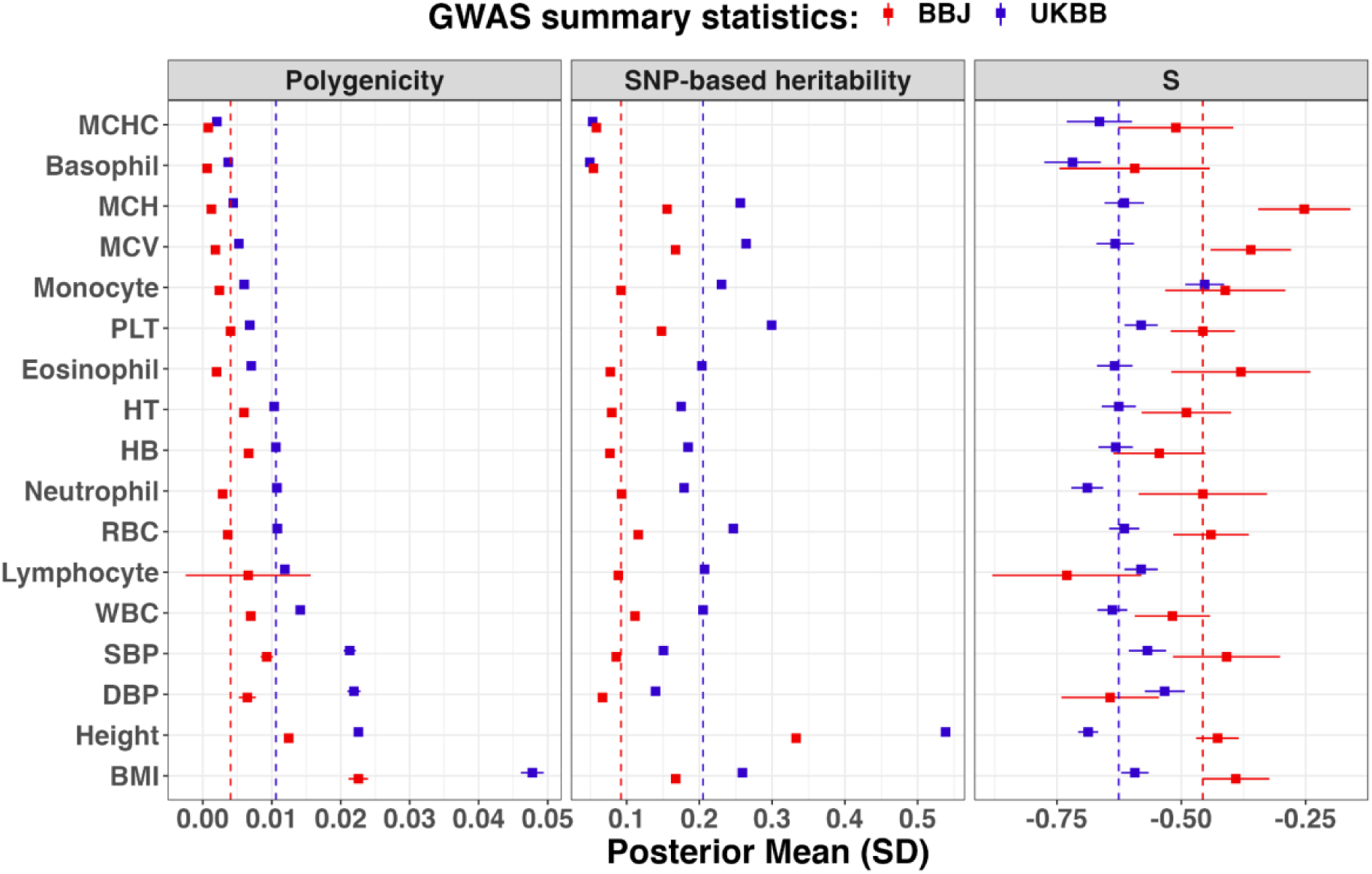
Genetic architecture of 17 studied traits between Biobank Japan (BBJ) and UK Biobank (UKBB). The error bar is the standard deviation of the corresponding estimate. The vertical dashed line was the median estimate. Full results are shown in **Table S4**. The phenotypes were ranked according to their polygenicity estimates using GWAS from UKBB, including: BMI (body mass index), Height, DBP (diastolic blood pressure), SBP (systolic blood pressure), WBC (white blood cell count), Lymphocyte (lymphocyte count), RBC (red blood cell count), Neutrophil (neutrophil count), HB (hemoglobin concentration), HT (hematocrit percentage), Eosinophil (eosinophil count), PLT (platelet count), Monocyte (monocyte count), MCV (mean corpuscular volume), MCH (mean corpuscular hemoglobin), Basophil (basophil count), MCHC (mean corpuscular hemoglobin concentration).

#### Multi-ancestry GWAS-derived PRS usually improves predictive performance compared to single-ancestry GWAS-derived PRS

We constructed PRS_single_ using the P+T and PRS-CS methods with GWAS from UKBB and BBJ, respectively. The GWAS sample sizes varied based on the number of Bin_Total_, which represented the total number of bins specific to each trait as shown in **Table S3.** Each bin consisted of 5,000 individuals randomly selected from the respective cohort. We found that employing target ancestry-matched GWAS, even with smaller sample sizes, yielded comparable accuracy to utilizing large-scale EUR GWAS but depended on PRS methodology and trait-specific genetic architecture (**Figure S6, Figure S7, Table S5** and **Supplementary Note 6**). We evaluated predictive accuracy by computing incremental *R*^2^ using linear regression, while accounting for the potential impact of covariates (**STAR Methods**).

For comparison, we developed PRS_multi_ using both P+T and PRS-CS, where we meta-analyzed single-ancestry GWAS from UKBB and BBJ. Similar to the simulation setup, we mimicked proportional ancestry composition in the multi-ancestry GWAS by meta-analyzing EUR GWAS in the UKBB with GWAS in the BBJ while varying number of bins (each bin of 5,000 individuals, UKBB bins ranging from 8 to 64 with an increment of 8, see **STAR Methods** and **Figure 1**). The ratio of EUR/EAS samples in the multi-ancestry GWAS varied from 64:1 to 8/Bin_Total_. Thus, 85% of the multi-ancestry GWAS had a higher proportion of EUR samples (>50% EUR). Consistent with our findings from the simulations, where we observed that the choice of LD reference panel had a limited impact on the predictive accuracy of more polygenic traits, we observed only a slight improvement of median *R*^2^ of 0.002 for P+T when employing a combined LD reference panel that was proportional to the ancestries represented in the multi-ancestry GWAS. We compared this result with PRS developed using a reference panel that was matched with the majority population of the discovery GWAS (**Figure S8 and Table S6**). Because the majority of PRS was constructed from GWAS predominantly composed of EUR individuals, we hereafter reported the results using 1KG-EUR as the LD reference.

In our analysis comprising 3,160 comparisons between single-ancestry PRS derived from UKBB GWAS (PRS_EUR_GWAS_) and multi-ancestry PRS (PRS_multi_), we observed encouraging results. Specifically, in the UK Biobank East-Asian population (UKBB-EAS), PRS_multi_ showed accuracy improvements in 99.7% and 92.4% of these comparisons when using P+T and PRS-CS, respectively (**Table S7 and Figure S9**). Accuracy increased with more EAS samples in the multi-ancestry GWAS (**Figure 4**). For example, when comparing PRS_multi_ with PRS_EUR_GWAS_ using P+T, the largest relative improvements in *R*^2^ were 80.9% (0.085 vs. 0.047) for platelet count (PLT), 152.2% (0.058 vs. 0.023) for BMI and 91.9% (0.071 vs. 0.037) for height. We observed these improvements when using multi-ancestry GWAS including EAS bins from BBJ, which were either concordant with or proximal to Bin_Total_, along with 64 EUR bins from UKBB. Similarly, the corresponding relative *R*^2^ improvements for these same three traits were 19.8% (0.0126 vs. 0.101), 50.0% (0.075 vs. 0.050) and 15.5% (0.097 vs. 0.084) when using PRS-CS. We did not consistently observe the upward trend for white blood cell count (WBC) with PRS-CS, which can be attributed to the lack of accuracy improvement with larger sample sizes of BBJ (**Figure S6**). We also found that P+T showed greater improvement compared to PRS-CS but worse accuracy overall, regardless of the number of bins from EUR GWAS; the median improvements in *R*^2^ across traits were 0.014 and 0.008, respectively. However, the upward trend in PRS accuracy was not consistently shown in the UKBB-EUR, particularly when using PRS-CS (**Figure S10 and Table S7**). This pattern aligned with our simulation results and previous reports that PRS accuracy for minority populations included in the multi-ancestry GWAS benefited more from adding more ancestry-matched individuals compared to other populations, including EUR populations^33^. We noted that the accuracy of PRS_multi_ remained largely unchanged or slightly decreased when the number of bins from BBJ was small (e.g., 1 or 2 bins), which was consistent with previous studies^6, 33^. In contrast to PRS derived from BBJ (PRS_Minor_GWAS_), we noted a diminishing trend in accuracy improvements of PRS_multi_ as the sample sizes of BBJ increased, especially for traits such as height, PLT, MCH and MCV (**Figure S11**). Furthermore, we observed greater variation in accuracy among traits from real data compared to simulations, which could be attributed to the smaller sample sizes and the more complicated genetic architecture.

**Figure 4.**
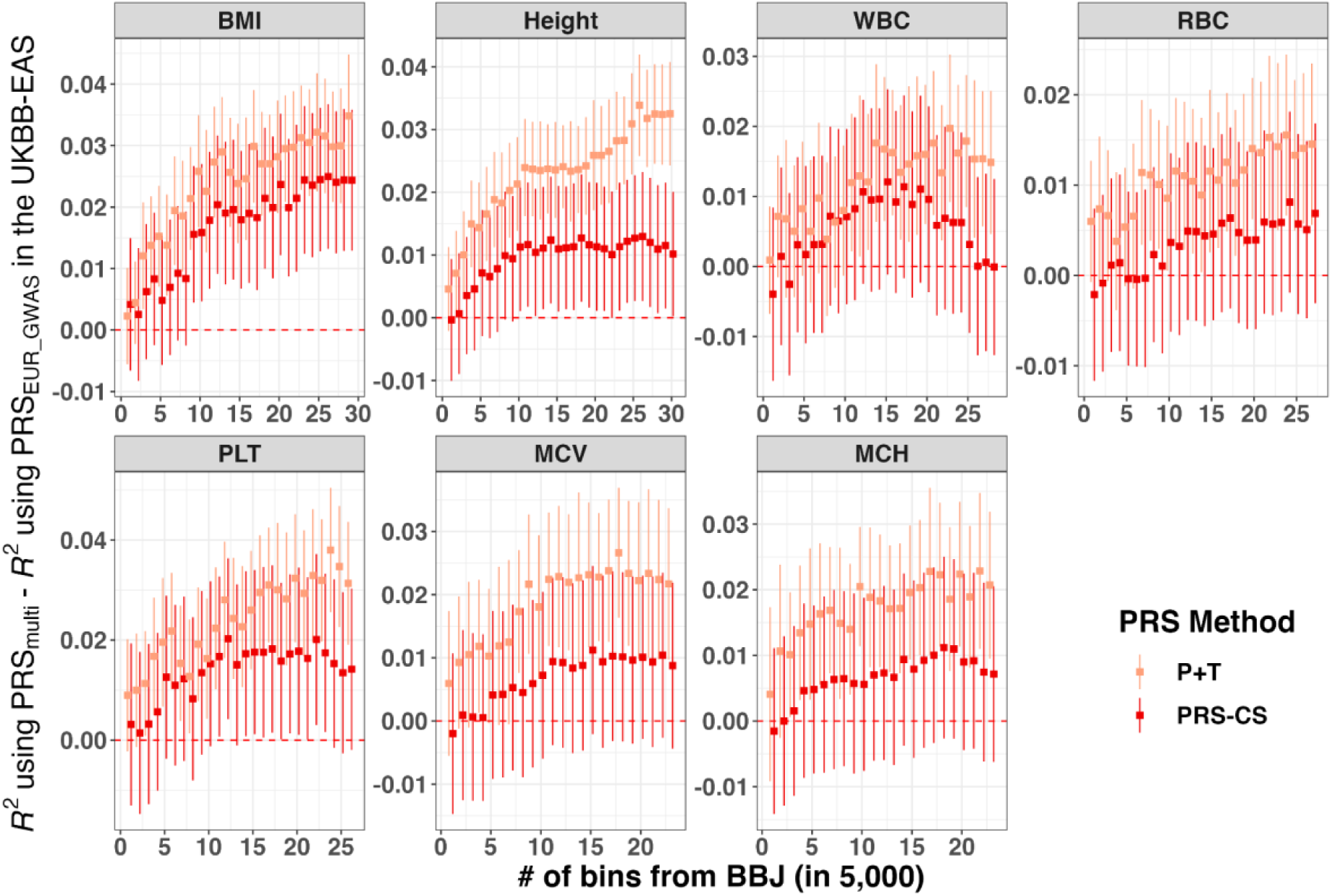
Accuracy improvement of PRS in the UK Biobank East-Asian population (UKBB-EAS) using multi-ancestry GWAS comprare to using European (EUR) GWAS for P+T and PRS-CS. The multi-ancestry GWAS were obtained by meta-analyzing EUR GWAS and EAS GWAS, with the EAS sample size from the Biobank Japan (BBJ) varying as indicated on the x-axis. For illustrative purposes, we present the results using 64 EUR bins, each containing 5,000 individuals, which were included in both EUR GWAS and multi-ancestry GWAS. PRS were constructed using P+T and PRS-CS and evaluated in the UKBB-EAS. The y-axis is the accuracy difference of PRS when using multi-ancestry GWAS (PRS_multi_) compared to using EUR GWAS (PRS_EUR_GWAS_). The error bars indicate the standard error of accuracy improvement. The red dashed line is y=0. We showed the results for 7 traits with SNP-based heritability > 0.1 in both BBJ and UKBB, and they were ranked by polygenicity estimates using UKBB (Figure 3). Full results are shown in **Table S7**.

#### PRS derived from meta-analyzed multi-ancestry GWAS versus weighted PRS from single-ancestry GWAS in understudied populations

In contrast to PRS_multi_, an alternative approach proposed in previous studies to enhance predictive accuracy in diverse populations is the linear combination of PRS derived from GWAS conducted on populations with different ancestries^34^. Here, we implemented this approach by developing a weighted PRS (**PRS_weighted_**) using P+T and PRS-CS. This combination involved linearly weighting PRS derived from single-ancestry GWAS conducted in the UKBB and BBJ. Additionally, we employed a more advanced Bayesian method called PRS-CSx^8^, which jointly models GWAS and LD information from multiple populations. Similarly, we constructed PRS_weighted_ using ancestry-specific posterior SNP effects. Furthermore, we developed PRS by integrating ancestry-specific posterior SNP effects using the inverse-variance weighted meta-analysis strategy, also referred to as PRS_multi_ (see **STAR Methods**).

Among the three PRS methods evaluated in the UKBB-EAS, PRS-CSx exhibited the highest performance, followed by PRS-CS and P+T. Specifically, for PRS_multi_, the corresponding median *R*^2^ values across traits were 0.051, 0.048 and 0.037, while for PRS_weighted_, they were 0.051, 0.045 and 0.021, respectively (**Figure 5, Table S8 and Table S9**). Notably, we observed that PRS_multi_ for BMI using PRS-CS yielded significantly better accuracy compared to PRS-CSx (median *R*^2^: 0.057 vs. 0.055, *p*-value < 2.2e-16, one-sided Wilcoxon signed-rank test). Out of the 3,160 comparisons between PRS_multi_ and PRS_weighted_ in the UKBB-EAS, 91.4% and 78.0% showed higher accuracy of PRS_multi_ when using P+T and PRS-CS, respectively, with median improvements in *R*^2^ of 0.011 (*p*-value < 2.2e-16) and 0.003 (*p*-value < 2.2e-16). Although we found better performance overall with PRS_multi_, we found that PRS_weighted_ significantly outperformed PRS_multi_ for PLT using P+T (median *R*^2^: 0.086 vs. 0.081, *p*-value < 2.2e-16) and for height using PRS-CS (median *R*^2^: 0.091 vs. 0.082, *p*-value = 2.6e-04). Contrary to trends observed with other methods, in 59.7% of the comparisons, PRS_weighted_ outperformed PRS_multi_ when using PRS-CSx, although we observed no significant accuracy difference across traits. However, PRS_weighted_ showed superior performance compared to PRS_multi_ (*p*-value < 0.05/17) for several traits, including MCV (median *R*^2^: 079 vs. 0.072), MCH (median *R*^2^: 0.079 vs. 0.073), Basophil (median *R*^2^: 0.010 vs. 0.007) and hemoglobin concentration (HB, median *R*^2^: 0.025 vs. 0.024).

**Figure 5.**
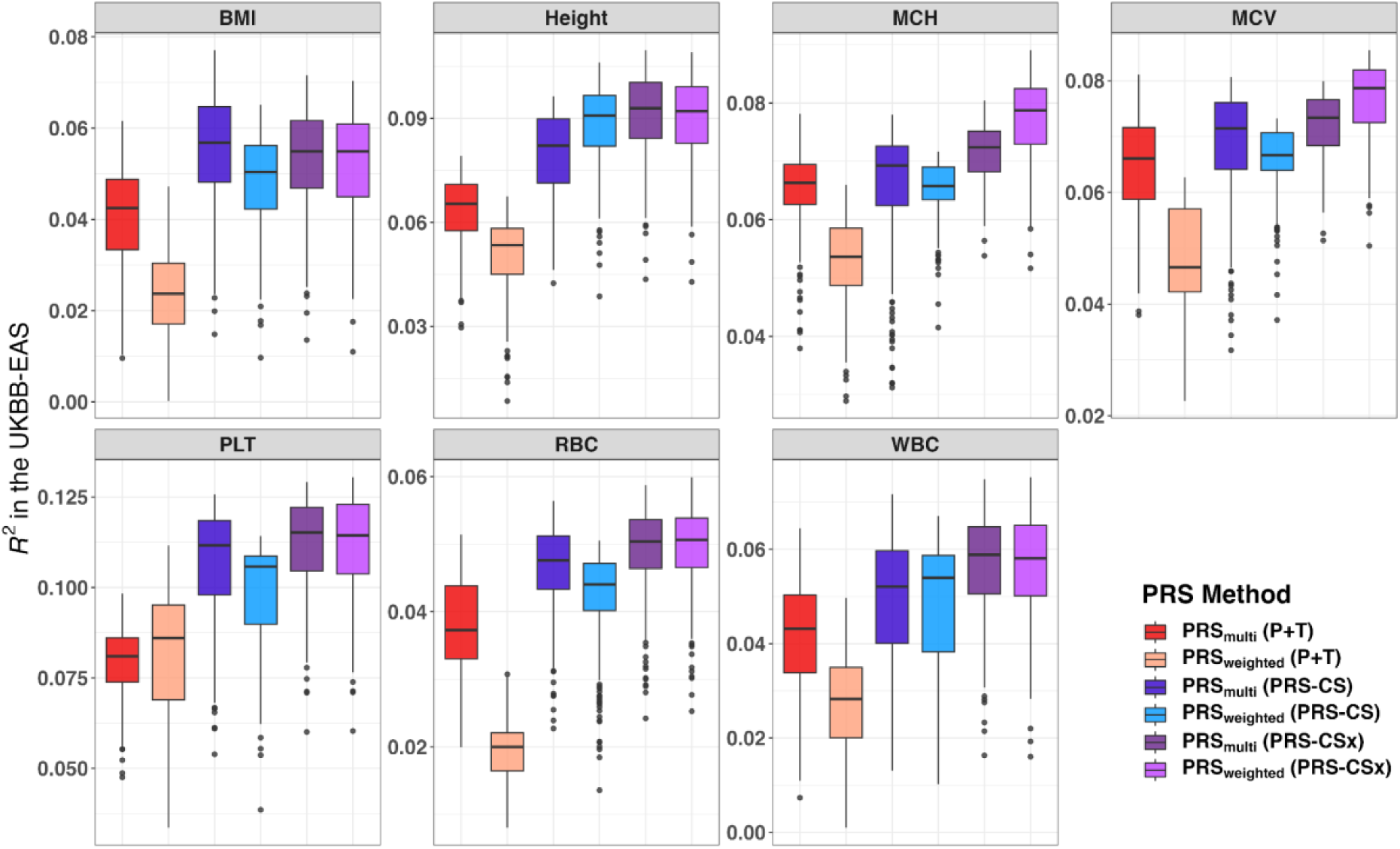
Predictive accuracy using different PRS methods in the UK Biobank East-Asian population (UKBB-EAS). PRS_mutli_ represents PRS derived from multi-ancestry GWAS, while PRS_weighted_ denotes PRS constructed from a weighted linear combination (see **STAR Methods** for details). PRS were constructed with three methods, including P+T, PRS-CS and PRS-CSx. We showed the results for 7 traits with SNP-based heritability > 0.1 in both Biobank Japan (BBJ) and UKBB. Traits were ranked by polygenicity estimates using UKBB (Figure 3). Boxes represent the first and third quartiles, with the whiskers extending to 1.5-fold the interquartile range. Full results are shown in **Table S8 and Table S9**.

Moreover, the extent of accuracy improvements using PRS_multi_, in contrast to PRS_weighted_, largely varied across traits and ancestry compositions. For example, when evaluating accuracy within the UKBB-EAS using P+T, we observed 3.25-fold increase in *R*^2^ with PRS_multi_ compared to PRS_weighted_ for monocyte count (monocyte, 0.065 vs. 0.020). This improvement was achieved with a bin ratio 56:15 for the discovery GWAS, consisting of 56 bins from UKBB and 15 bins from BBJ. Similarly, using a bin ratio of 40:25, we achieved a 4-fold increase in *R*^2^ for DBP (0.048 vs. 0.012) with PRS_multi_ compared to PRS_weighted_. When developing PRS_multi_ using PRS-CS, we observed notable relative improvements in *R*^2^ when compared to PRS_weighted_, specifically a 24.7% increase for PLT (0.091 vs. 0.073) with bin ratio of 24:1, and a 57.1% increase for lymphocyte (0.044 vs. 0.028) with a bin ratio of 16:1. Additionally, we found that PRS-CSx showed better performance in comparison to PRS-CS, especially when EUR GWAS was smaller or Minor GWAS was larger.

However, such improvements were less pronounced with large-scale EUR GWAS or small Minor GWAS (**Figure S12**). While sharing ancestry-specific GWAS summary statistics is highly beneficial for determining optimal approaches, our findings highlight the value of pragmatic approaches that directly construct PRS from large-scale meta-analyzed multi-ancestry GWAS. Such studies are often more accessible than ancestry-specific GWAS summary statistics.

### PRS derived from local ancestry-informed GWAS can improve accuracy for some less polygenic traits

We next conducted a comparative analysis to evaluate the optimal PRS approaches for admixed populations, utilizing local ancestry-informed GWAS. Specifically, we used Tractor^19^ to perform GWAS in AFR tracts within admixed AFR-EUR individuals, referred to as **AFR_Tractor_**. This approach enabled us to construct ancestry-specific PRS across 17 traits in the understudied AFR population. We developed PRS using both P+T and PRS-CS, and subsequently compared the accuracies of PRS derived from AFR_Tractor_ with those derived from large-scale EUR GWAS performed with standard linear regression (**EUR_Standard_**). To maximize discovery sample size, we also developed PRS_weighted_ by combining EUR_Standard_-derived PRS and AFR_Tractor_-derived PRS through linear weighting; we compared its performance to PRS derived from multi-ancestry meta-analyzed GWAS (referred to as **Meta_Standard_**, see **STAR Methods**).

Local ancestry-informed ancestry-specific GWAS had a much smaller sample size relative to the EUR-inclusive GWAS, as is typical for GWAS of underrepresented populations. As expected, we did not observe significant predictive accuracy of AFR_Tractor_-derived PRS for most traits such as height and BMI (**Figure 6 and Table S10**). However, we observed notable improvements for 5 traits, including WBC, neutrophil count (neutrophil), MCV, MCH and MCHC, where AFR_Tractor_-derived PRS achieved significantly higher *R*^2^ compared to EUR_Standardr_-derived PRS when using P+T (0.040 vs. 0.007, one-sided paired *t*-test, *p*-value = 0.038), despite a much larger sample size for EUR_standard_. This improvement might be attributed to the presence of large-effect AFR-enriched variants, particularly for MCV, MCH and MCHC, which are effectively captured by Tractor GWAS^6, 19^. Consistent with our previous findings, P+T generally outperformed PRS-CS for these traits characterized by much sparser genetic architectures, with a mean *R*^2^ of 0.040 compared to 0.022. Given that heritability bounds predictive accuracy, which can vary among populations and contexts, we also compared SNP-based heritability estimates between the AFR and EUR populations in the Pan-UK Biobank Project (https://pan.ukbb.broadinstitute.org/docs/heritability/index.html). In line with our PRS accuracy results, we observed higher estimates of SNP-based heritability for WBC (*h*^2^ = 0.41, SE = 0.19 vs. *h*^2^ = 0.17, SE = 0.01), neutrophil (*h*^2^ = 0.44, SE = 0.26 vs. *h*^2^ = 0.15, SE = 0.01), and MCHC (*h*^2^ = 0.15, SE = 0.11 vs. *h*^2^ = 0.06, SE = 0.01) in the AFR population compared to the EUR population. However, these differences did not reach statistical significance, which can be attributed to the large standard errors resulting from the limited small sample size of AFR population and the sparser genetic architectures, leading to less stable heritability estimates using LDSC.

**Figure 6.**
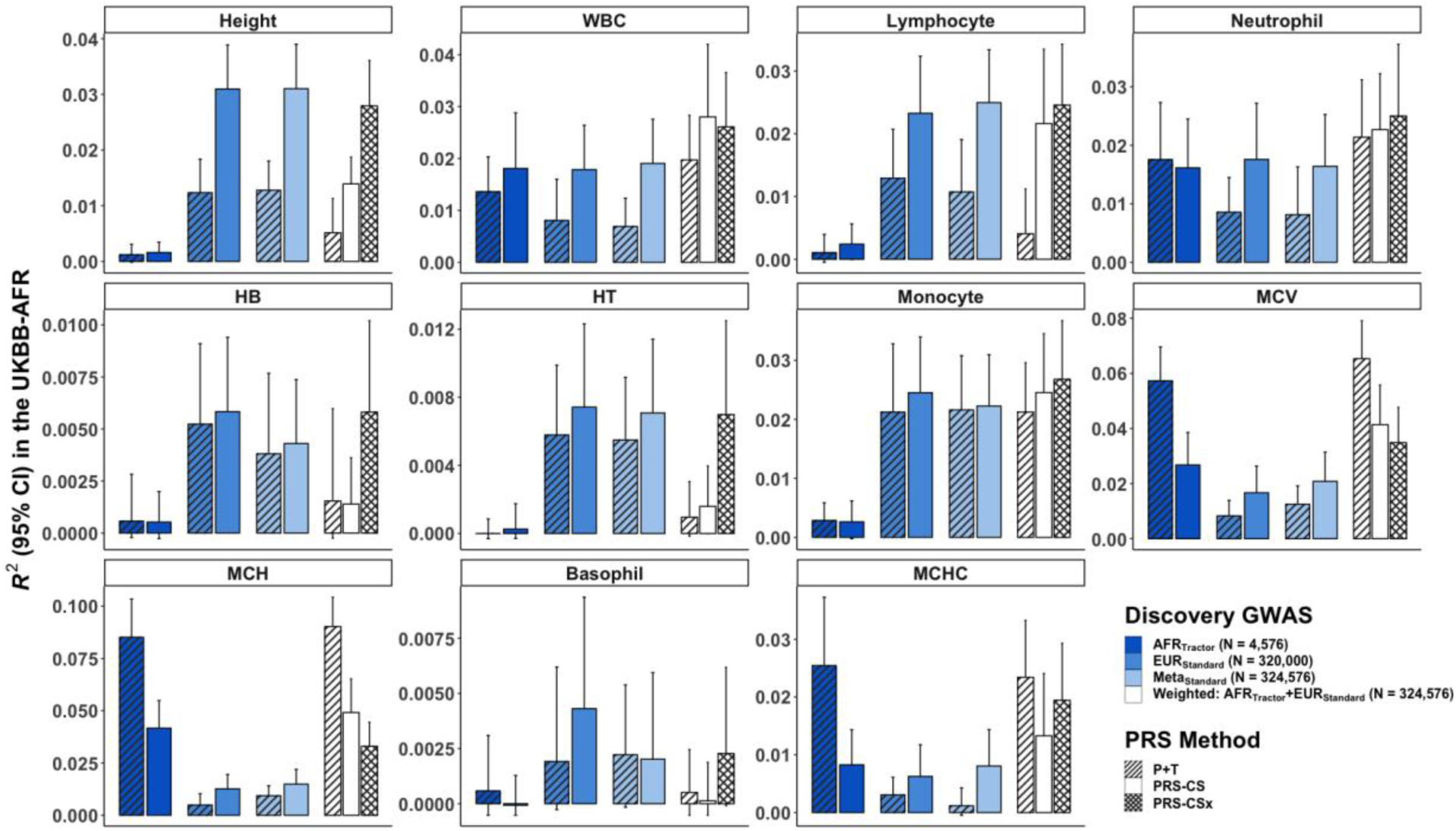
Accuracy of PRS derived from local-ancestry informed GWAS versus other discovery GWAS in the UK Biobank African population (UKBB-AFR) We evaluated PRS performance in the UKBB-AFR by utilizing various methods on different discovery GWAS. Specifically, AFR_Tractor_ denotes the AFR-specific GWAS performed using Tractor on the UKBB admixed African-European individuals. EUR_Standard_ refers to standard GWAS performed on the European (EUR) population in the UKBB. Meta_Standard_ is the meta-analysis performed on AFR_Tractor_ and EUR_Standard_. Furthermore, we constructed a weighted PRS by combining PRS generated from AFR_Tractor_ and EUR_Standard_ through a linear weighted approach. The figure shows the results for traits with SNP-based heritability > 0.1 in the UKBB-AFR. Full results are shown in **Table S10**.

The best local ancestry-informed PRS approach that we evaluated for the 5 less polygenic traits with large ancestry-specific effects was a weighted linear regression approach. This approach combined PRS derived from AFR_Tractor_ and EUR_Standard_ using linear regression and outperformed predictive accuracy compared to using Meta_Standard_-derived PRS. This finding aligns with our earlier observations, where PRS_weighted_ outperformed PRS_multi_ for traits with large effect ancestry-enriched variants, while PRS_multi_ exhibited superior overall performance for traits lacking such variants. Specifically, the mean accuracies of PRS_weighted_ using P+T, PRS-CS and PRS-CSx for those 5 traits were 0.044, 0.031, and 0.028, respectively, with no significant differences observed among the three PRS methods. The mean accuracies of Meta_Standard_ were 0.016 and 0.008 using PRS-CS and P+T, respectively. Additionally, we did not observe significant accuracy differences between PRS derived from GWAS conducted using standard linear regression in admixed populations and AFR_Tractor_-derived PRS **(Table S10**). It is worth noting that the effective sample size of local ancestry-informed GWAS is approximately 20% smaller due to the reduction from deconvolving ancestral tracts. Moreover, PRS derived from traditional GWAS in admixed populations necessitate an in-sample LD reference panel. In contrast, local ancestry-informed GWAS-based PRS, as shown in this study, can leverage external LD reference panels, eliminating the need for direct access to individual-level genotypes of admixed populations.

## Discussion

In this study, we extensively evaluated PRS performance through a combination of simulation and empirical analyses to explore the impact of various factors on PRS predictive accuracy and generalizability across populations. We demonstrated that increasing genetic diversity of discovery GWAS improved predictive accuracy in understudied populations. The extent of improvement was influenced by factors such as sample size ratios between EUR GWAS and Minor GWAS, genetic architecture, PRS methodology, and LD reference panels. Among those factors, between-ancestry genetic architecture differences, such as ancestry-enriched variants with large effects, affected accuracy improvement more than other factors. While leveraging large-scale EUR GWAS continues to benefit PRS accuracy given the current scale of understudied populations, we may not expect accuracy improvement when meta-analyzing extremely small Minor GWAS^24^.

Our study also revealed that directly meta-analyzing datasets from diverse ancestral groups could yield greater accuracy improvements than linearly combining PRS through an optimized weighting strategy, especially for P+T. Such improvements from meta-analyzed GWAS supports the common implicit assumption that causal variants are shared between ancestries. Consistent with this assumption, when smaller target populations lack representation, leveraging genetic information from a different population with larger sample sizes improves PRS accuracy, even when it is ancestrally diverged. Notably, when employing the more sophisticated genome-wide PRS method, PRS-CSx, accuracy differences between PRS_multi_ and PRS_weighted_ were marginal. Moreover, PRS-CSx generally outperformed PRS-CS, with the exception of BMI. The improvement was most pronounced for traits with ancestry-specific variants, such as MCV and MCH.

We have comprehensively evaluated characteristics that impact PRS performance, including in recently admixed populations. We have shown the advantage of leveraging GWAS in admixed populations by accounting for local ancestry, which could improve PRS predictive performance in understudied populations even without direct access to individual genotypes of admixed populations. Specifically, we found that PRS_weighted_ consistently outperformed PRS_multi_ for traits with ancestry-enriched variants. However, the sample size of admixed individuals here was relatively small, and we anticipate that future analyses incorporating larger datasets, such as the *All of Us* Research Program, will provide further insights into optimal PRS strategies for improved accuracy and generalizability using PRS derived from local ancestry-informed GWAS.

While previous studies have shown the advantages of leveraging increased genetic diversity to improve PRS accuracy in global populations^7, 35^, most have used GWAS with primarily European ancestry. Here, we have provided additional best practices for developing PRS for understudied populations using diverse discovery cohorts, particularly when GWAS encompass different ancestry compositions across various trait genetic architectures (**Figure 7**). Our recommendations primarily revolve around general guidelines for constructing PRS_single_ and PRS_multi_ (or PRS_weighted_), depending on factors examined in this study (**Figure S13**).

**Figure 7.**
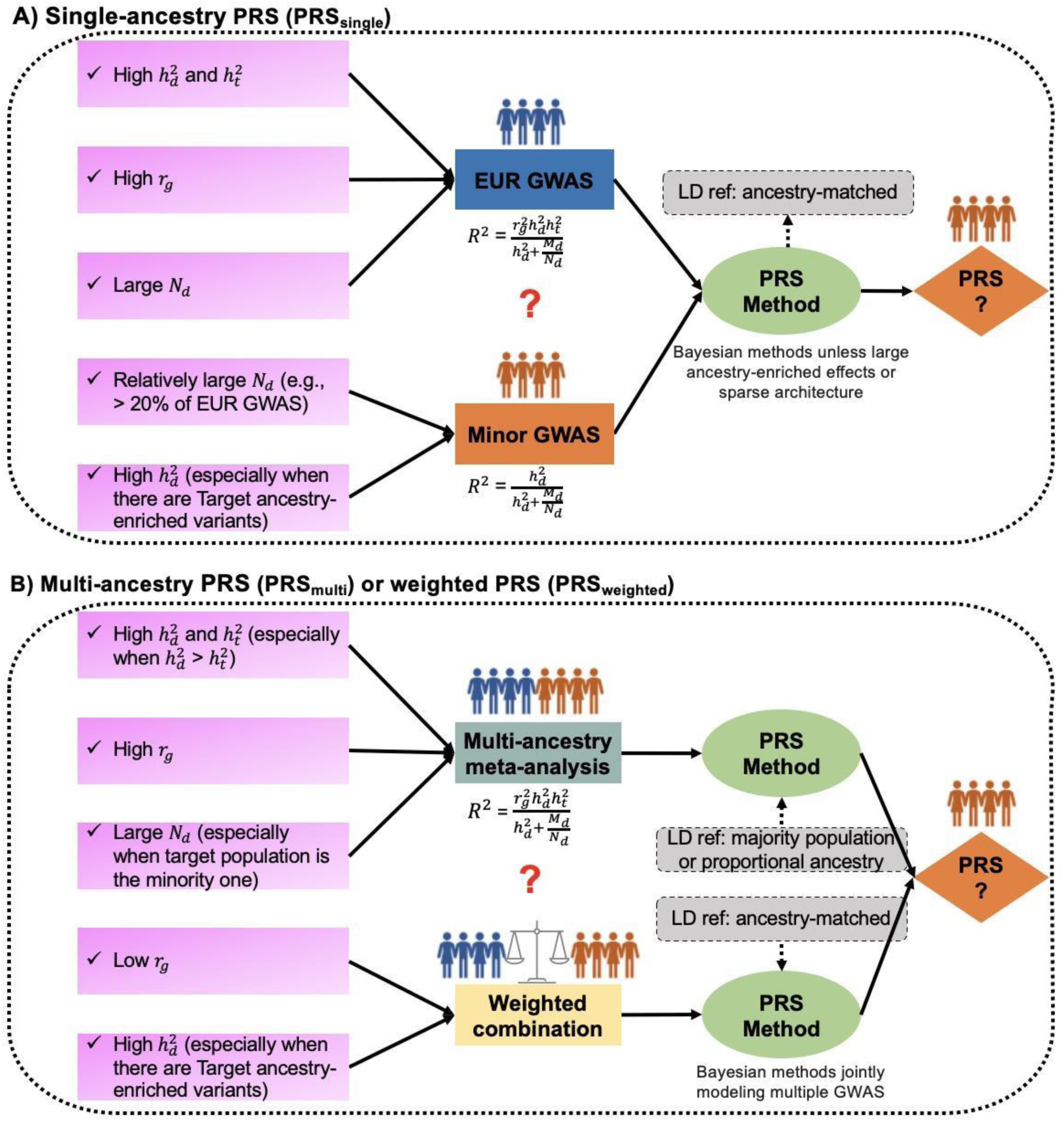
General practices for developing PRS using different discovery GWAS. We summarized the general practice for developing PRS A) using single-ancestry GWAS (PRS_single_); and B) using GWAS from multiple ancestries (PRS_multi_ or PRS_weighted_). Abbreviations: Cross-ancestry genetic correlation (*r_g_*), SNP-based heritability in discovery 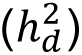 and target populations 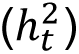, discovery GWAS sample size (*N_d_*) and the number of genome-wide independent segments in the discovery population (*M_d_*).

First, in the development of PRS_single_, we employed a theoretical equation^36^ to enhance the selection of input GWAS (**Supplementary Note 7),** as a function of the cross-ancestry genetic correlation, SNP-based heritability in discovery and target populations, discovery GWAS sample size, and the number of genome-wide independent segments in the discovery population^36^. For traits with relatively low *r_g_* and a sizable ancestry-matched GWAS (e.g., > 20-40% of EUR GWAS), such as BMI and height, PRS accuracy in the target population improves when ancestry-matched GWAS are utilized. On the other hand, for traits with high *r_g_* and SNP-based heritability, we expect larger-scale EUR GWAS to outperform smaller-scale ancestry-matched GWAS. However, it is important to consider the characteristics of the target cohort and phenotype precision. Additionally, we expect Bayesian methods tailored to trait-specific genetic architecture to outperform classic P+T methods. However, this superior performance may not hold true for traits that exhibit large-effect ancestry-enriched variants or with a very sparse genetic architecture, which are attributes typically informed by prior knowledge or information gleaned from literature and public resources^35, 37–39^. To enhance accuracy in such scenarios, we recommend employing a grid-search approach with a finer-scale adjustment of the hyper-parameters in Bayesian methods.

Second, in comparison to PRS_single_ derived from large-scale EUR GWAS, we recommend using PRS_multi_, unless the target ancestry-matched GWAS is extremely small (<10,000). PRS_multi_ is generally preferred for traits with high *r_g_* SNP-based heritability, and large sample sizes. We find increasing evidence supporting the notion that the effects of most common variants are shared between ancestries, indicating a high *r_g_* for most traits^9, 11^. However, estimates of *r_g_* can be affected by phenotypic and environmental heterogeneity across populations^10, 40^. When constructing PRS_multi_ using summary-level based methods such as P+T and PRS-CS, researchers should carefully consider which LD reference panel best approximates the LD structure between SNPs while being most readily accessible. We have shown that when EUR remains the majority population in the discovery GWAS, using the EUR-based reference panel effectively approximates the LD of discovery GWAS, consistent with our previous findings^7^.

Third, our findings indicate the advantages of PRS_multi_ compared to PRS_weighted_, particularly when employing P+T and PRS-CS. However, there are some notable exceptions, such as the higher accuracy observed when using PRS_weighted_ with PRS-CS for traits with low *r_g_*, such as height. Furthermore, when incorporating local ancestry-informed GWAS and large-scale EUR GWAS, PRS_weighted_ outperformed PRS_multi_ for traits with AFR-enriched variants, such as WBC and MCHC, in the UKBB-AFR. On the other hand, we note that the accuracy of PRS_multi_ could be more affected by the choice of LD reference panel, while PRS_weighted_ was not limited in this regard due to its easy accessibility of external ancestry-matched reference panels. PRS-CSx, which accounts for variations in allele frequencies and LD structures across ancestries, is recommended when ancestry-specific GWAS from multiple populations are available, especially with considerable sample sizes (e.g., > 25,000∼50,000) in the Minor GWAS. These results highlight the importance of making ancestry-specific GWAS summary statistics publicly available.

In summary, there is no one-size-fits all approach for constructing PRS, as the optimal approach depends on genetic architecture, ancestry composition, statistical power, and other factors. These factors can be complex, particularly as a deluge of methods are being developed to address the PRS generalizability problem. To inform optimal approaches across a wide range of scenarios, we have distilled the results of extensive simulations and empirical analyses across trait genetic architectures, ancestries, and methods into a set of guidelines from parameters that are typically evaluated at the outset of a genetic study.

### Limitations of the study

We acknowledge some limitations and future directions in our study. First, we focused on common variants in different populations, while population-enriched variants have lower frequencies and larger effect sizes in some populations. The role of such variants in polygenic prediction are worth exploring across phenotypes when there are sufficient sample sizes for different ancestral populations. Second, as we used external LD reference panels for PRS construction, PRS performance decreases with LD mismatch between the discovery population and LD reference panel, especially using multi-ancestry GWAS. While we show that LD reference panel differences have a relatively modest effect on PRS accuracy, they have a much larger effect on fine-mapping^41^, so future efforts are warranted to share in-sample LD without direct access to individual-level genotypes, especially for large consortia with numerous and diverse cohorts. Alternatively, developing more sophisticated individual-level PRS methods that preserve privacy and are scalable to current biobank-scale data is also promising. Third, while our primary focus pertains to quantitative phenotypes characterized by diverse genetic architectures, we expect our findings can be broadly applied to binary traits, as we have investigated previously^7^. However, binary phenotypes introduce additional complexities due to factors such as variable case/control ratios, phenotype definitions, environmental differences, and smaller effective sample sizes or lower statistical power. Fourth, while we have provided theoretical expectations of cross-ancestry prediction, the reliability of parameter estimates such as cross-ancestry genetic correlation and the effective number of independent genome-wide segments poses significant challenges, particularly in the context of multi-ancestry GWAS with highly imbalanced sample sizes. Finally, it is important to acknowledge that our study focused on selected methods, which consistently exhibit similar trends^42^. Although we anticipate that our findings are broadly applicable to alternative methods, such as XPASS^43^ and XP-BLUP^42^, further research is needed to explore the generalizability of our findings to other polygenic prediction approaches. Despite the limitations, our study highlights the advantages of leveraging the increasing diversity of current genomics studies to improve polygenic prediction across populations. We emphasize the necessity of diversifying not only the ancestry but also phenotypic spectrum when collecting genomic data from global populations, which will contribute to achieve a more equitable and effective use of PRS for traits with varying genetic architectures.

## Acknowledgements

A.R.M. and Y.W. were supported by funding from the National Institutes of Health (K99/R00MH117229 to A.R.M.) as well as funding from European Union’s Horizon 2020 research and innovation program under grant agreement 101016775. A.R.M. was also supported by funding from U01HG011719. H.H. acknowledges support from NIDDK K01DK114379, NIDDK 1R01DK129364, NIMH U01MH109539, the Zhengxu and Ying He Foundation, and the Stanley Center for Psychiatric Research. E.G.A. is supported by K01MH121659 from NIMH, the Caroline Wiess Law Fund for Research in Molecular Medicine, and the ARCO Foundation Young Teacher-Investigator Fund at Baylor College of Medicine. P.T. is supported by funding from NIH/NIA (R00 AG062787) and by the Good Ventures Foundation (010623-00001).

## Author Contributions

Conceptualization: Y.W., A.R.M., E.G.A., M.Kanai.

Formal analysis: Y.W., M.Kanai., T.T., M.Kamariza., K.Y., W.Z., P.T.

Writing – Original Draft: Y.W., A.R.M., E.G.A., M.Kanai., T.T.

Writing – Review & Editing: Y.W., M.Kanai., T.T., M.Kamariza, K.T., K.Y., W.Z., Y.O., H.H., P.T., K.T., E.G.A., A.R.M.

## Declaration of interests

H.H. received consultancy fees from Ono Pharmaceutical and honorarium from Xian Janssen Pharmaceutical. All other authors declare no competing interests.

## STAR Methods

### Resources Availability

#### Lead Contact

Further information and requests for resources and reagents should be directed to and will be fulfilled by the lead contact, Ying Wang.

#### Materials Availability

This study did not generate new unique reagents.

#### Data and code availability

1000 Genome Phase 3 data can be accessed at ftp://ftp.1000genomes.ebi.ac.uk/vol1/ftp/data_collections/1000_genomes_project/data. We used UK Biobank data via application 31063. The software used in this study can be found at: Plink (https://www.cog-genomics.org/plink/), PRS-CS (https://github.com/getian107/PRScs), PRS-CSx (https://github.com/getian107/PRScsx), Tractor (https://github.com/Atkinson-Lab/Tractor), HapGen2 (https://mathgen.stats.ox.ac.uk/genetics_software/hapgen/hapgen2.html) and SBayesS/GCTB (https://cnsgenomics.com/software/gctb/). The Pan UK Biobank Project can be accessed at: Pan-UK Biobank Project https://pan.ukbb.broadinstitute.org. The codes used in this study have been deposited to https://github.com/ywangleo/multi-ancestry-PRS.

## Methods Details

### Simulations

#### Simulated genotypes in three populations

To explore the potential improvement of predictive accuracy within an underrepresented target ancestry through the inclusion of additional samples included in the multi-ancestry discovery GWAS, we simulated genotypes of chromosome 22 for 560,000 individuals in each population including European ancestry (EUR), East Asian ancestry (EAS) and African ancestry (AFR) using the software HapGen2 v2.1.2^44^. We used the haplotypes from 1000 Genome Project (1KG, Phase 3)^45^ as the sample pool. We excluded Americans of African Ancestry in SW USA and African Caribbeans in Barbados from the AFR samples due to their high degree of recent admixture. We used default parameters in HapGen2 with effective sample sizes of 11,375, 12,239 and 17,380 for EUR, EAS and AFR, respectively^44^. After simulating the genotypes on chromosome 22, we ran analyses with a total of 87,938 overlapping SNPs across the three ancestries which passed quality control filters: minor allele frequency (MAF) > 0.01, Hardy-Weinberg Equilibrium (HWE) *p*-value > 10*^−^*^6^ and genotype missingness rates across individuals < 0.05. We then removed 2nd-degree related individuals using the software KING^46^, resulting in 534,352, 533,996 and 537,498 unrelated individuals from EUR, EAS and AFR, separately. We randomly sampled 10,000 and 520,000 individuals from each ancestry as the withheld target population and discovery population, respectively.

#### Simulated phenotypes with varying trait genetic architecture

For the sake of simplicity, we assumed that causal variants are shared across populations and their effect sizes are perfectly correlated (cross-ancestry genetic correlation, *r_g_* = 1) in our initial simulations. The pairwise *r_g_* among *K* populations is represented by a *K* ∗ *K* matrix, denoted as **R,** where the off-diagonal elements of **R** had the value of *r_g_* and diagonal elements of **R** were set to 1. In our study, *K* was equal to 3, indicating the number of populations considered. We simulated phenotypes based on the simple additive model: *y* = *g* + *e*, where 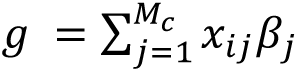*. *M_c_* is the number of causal variants, *x_ij_* is the genotype coded as 0, 1, or 2 for the *j*th SNP in the *i*th population. The effect size of *j*th SNP across *K* populations is drawn from a multivariate normal distribution, *β*∼*MNN*(*0*, *Σ*)*, where for the *K* ∗ *K* variance-covariance matrix, *Σ* the diagonal and off-diagonal elements were 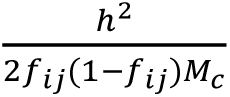 and 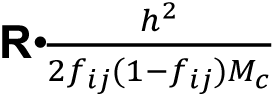, respectively. We denoted *f_ij_* as the MAF of *j*th SNP in the *i*th population and *h*^2^ as the trait heritability. We simulated the environmental effects to follow a normal distribution with 0 mean and 1 *− h*^2^ variance, *e* ∼ *N*(0, 1 *−h*^2^). We simulated different levels of heritability for chromosome 22 (*h*^2^ *=* 0.03 and 0.05). Additionally, we randomly sampled various numbers of causal variants (*M_c_* = 100, 500, and 1000) from all the 87,938 SNPs. As a result, we defined a total of 6 distinct simulation scenarios that encompass a realistic spectrum of polygenicity, ranging from ∼0.1% to ∼1% of causal variants. To assess the impact of *r_g_* on PRS performance, we expanded our simulation study by considering two scenarios. These scenarios aimed to capture different levels of per-variant variance explained. In scenario 1 characterized by *M_c_* = 100 and *h*^2^ *=* 0.05, the per-variant variance explained was higher. Conversely, scenario 2 involved *M_c_* = 1000 and *h*^2^ *=* 0.03, resulting in a lower per-variant variance explained. For each scenario, we varied the values of *r_g_* to 0.6 and 0.8, respectively.

#### Downsampling and meta-analyzed GWAS in simulations

To provide the requisite discovery data for constructing PRS, we proceeded to perform GWAS on the simulated phenotypes. Specifically, we split the discovery population, which consisted of 520,000 unrelated individuals, into 52 evenly distributed bins, each comprising 10,000 individuals (denoted as Bin_1_, Bin_2_,…, Bin_total_). Subsequently, we ran GWAS on each of those 52 bins independently within the three populations, using simple linear regression implemented in PLINK v2.0^47^. We excluded the causal variants when running GWAS to mimic the phenomenon of imperfect tagging. We then employed an iterative process of meta-analysis, employing the inverse-variance weighted method using METAL^48^, gradually incorporating a varying number of bins. Specifically, we commenced the meta-analysis with Bin_1_+Bin_2_, subsequently progressing to Bin_1_+Bin_2_+Bin_3_, and so forth, until we encompassed the complete set of bins (Bin_1_+Bin_2_+Bin_3_+…+Bin_tota_) for each population.

To simulate a scenario resembling a meta-analysis involving multiple ancestries with varying proportions, we opted for an arbitrary selection of subsets from EUR GWAS. Specifically, we chose a range of bins, from 4 to 52 bins, with increments of 4. Subsequently, we systematically incorporated different numbers of bins, spanning from 1 to 52, from EAS and AFR populations into the EUR GWAS dataset via meta-analysis. The meta-analysis was conducted utilizing the inverse-variance weighted fixed effects model implemented in the METAL software. This iterative process allowed us to achieve a range of sample size ratios between EUR and EAS as well as EUR and AFR, encompassing ratios from 52:1 to 4:52, in the meta-analyzed multi-ancestry GWAS (referred to as **Meta**). The simulation configuration is visually depicted in **Figure 1**.

#### Pruning and Thresholding (P+T) in simulations

We employed PLINK v1.90 to clump quasi-independent SNPs within 500Kb windows, utilizing a LD threshold of *r*^2^ < 0.1. To explore the impact of various LD reference panels on predictive accuracy of PRS, we used a total of four different LD reference panels: one for single-ancestry and three for multi-ancestry GWAS, with consideration to the ancestry composition of the discovery GWAS and the target population.

For the single-ancestry GWAS, we used a LD reference panel consisting of 10,000 individuals from the target population that were matched to the ancestry of the discovery GWAS. In the case of multi-ancestry GWAS, we used three LD reference panels. These panels included two composed of a single ancestry that did not mirror the ancestral makeup of the discovery GWAS. Specifically, one panel comprised 10,000 withheld EUR individuals, while the other panel encompassed individuals from understudied populations, either 10,000 EAS or 10,000 AFR individuals, consistent with the minority population represented in the discovery GWAS. The third LD reference panel consisted of individuals from different ancestries in proportions proportional to the discovery GWAS, amounting to a total of 10,000 samples.

We calculated PRS in the target population using 8 different *p*-value thresholds: 5 × 10^-8^, 1 × 10^-6^, 1× 10^-4^, 1 × 10^-3^, 0.01, 0.05, 0.1, and 1. We denoted PRS constructed from single-ancestry GWAS as single-ancestry PRS (PRS_single_) and those from meta-analyzed multi-ancestry GWAS as multi-ancestry PRS (PRS_multi_). We calculated the predictive accuracy as the variance explained by the PRS (*R*^2^) through linear regression: *y* ∼ PRS and computed corresponding 95% confidence intervals (CIs) through bootstrap. To identify the optimal *p*-value threshold associated with the highest predictive accuracy, we evenly divided the target population into a test cohort and a validation cohort. The *p*-value threshold was optimized through a process of hyperparameter tuning in the validation cohort, and subsequently, the accuracy of the model was assessed using the test cohort.

### Empirical analysis of 17 quantitative traits in the UK Biobank (UKBB) and Biobank Japan (BBJ)

We further explored how the findings from simulations generalized in real data using 17 quantitative traits shared between UKBB and BBJ, including anthropometric traits (BMI and height) and blood panel traits studied previously (**Table S3**)^32^. The selection of these traits was motivated by their widespread availability within biobanks and their substantial statistical power, attributable to their quantitative properties.

#### Datasets and Quality Control (QC)

##### UK Biobank (UKBB)

The details of assigning ancestry for each individual in the UKBB are described in the Pan-UK Biobank Project (Pan UKBB: https://pan.ukbb.broadinstitute.org/). Briefly, a random forest classifier trained on reference data from 1KG and Human Genome Diversity Project (HGDP)^49^ was used to classify cohort individuals under continental population labels based on the top 6 principal components (PCs). In this study, we used a total of 361,144 and 2,684 unrelated EUR and EAS participants, respectively. We obtained unrelated individuals through running hl.maximal_independent_set using Hail (https://hail.is/). Specifically, within each population, we ran PC-Relate^50^ with k=10 and min_individual_maf=0.05. We used the individuals assigned EAS ancestry as the target dataset. For EUR samples, we first randomly retained 5,000 individuals with complete phenotype information for all 17 studied phenotypes as the target population. Subsequently, we split the remaining individuals into evenly distributed bins, each containing 5,000 individuals, for each phenotype. The number of total bins for each studied phenotype ranged from 68 to 71, depending on phenotype missingness (**Table S3**). The bins were labeled sequentially from 1 to the total number of bins, following the same procedure as described in our simulations.

##### BioBank Japan (BBJ)

BBJ is a multi-institutional hospital-based biobank which has recruited approximately 200,000 participants from 12 medical institutions in Japan between fiscal years 2003 and 2007^27^. Written informed consents were obtained from all the participants, as approved by the ethics committees of the RIKEN Center for Integrative Medical Sciences, and the Institute of Medical Sciences, the University of Tokyo. The participants were genotyped using either (i) the Illumina HumanOmniExpressExome BeadChip or (ii) a combination of the Illumina HumanOmniExpress and HumanExome BeadChips. The genotypes were then prephased using Eagle^51^ and imputed using Minimac3^52^ with a reference panel that consists of 1KG samples (N = 2,504) and whole-genome sequencing (WGS) data of Japanese individuals (N = 1,037)^53^. Standard quality controls of participants and genotypes were applied as described elsewhere^53^. Briefly, we excluded samples with low call rates (< 98%), closely related individuals (PLINK PI_HAT > 0.175), or non-Japanese outliers based on the principal component analysis (PCA). We then excluded genotyped variants with call rate < 98%, HWE P-value < 1.0 × 10^−6^, number of heterozygotes < 5, or low concordance rate (< 99.5%) with WGS for a subset of individuals (N = 939). Phenotypes were retrieved from medical records and prepared as described previously^54^.

##### 1000 Genomes Project Phase 3 (1KG)

We used 1KG phase 3 data as LD reference panels in this study. Specifically, we kept 495 unrelated EUR, 498 unrelated EAS, and 484 unrelated AFR individuals from 1KG. The AFR individuals were solely utilized for analyses pertaining to recently admixed populations.

##### Quality Controls

The imputation strategies for UKBB and BBJ have been described in detail elsewhere^55, 56^. After imputation, we first excluded ambiguous variants (e.g., A/T and C/G) and further filtered to keep those variants with imputation INFO score > 0.3, MAF > 0.01, HWE p-value > 10^-6^, and genotyping missing rates across individuals < 0.05. Consequently, approximately 8.6 million and 6.6 million SNPs were retained for the UKBB and BBJ, respectively. For our analyses, we exclusively utilized SNPs that passed these quality control measures, resulting in approximately 3.6 million SNPs that were shared among both biobanks and 1KG.

#### PRS construction for 17 traits in empirical analysis

##### Discovery GWAS

All phenotypes were curated and transformed to be normally distributed as described previously^32^. Subsequently, we performed GWAS on the rank normalized phenotypes using simple linear regression implemented in PLINK v2.0. We included age, sex, age^2^, age × sex, age^2^ × sex, and the first 20 PCs as the covariates. In line with the GWAS strategy outlined in the *Simulations* section, we initially performed GWAS within individual bins and then engaged in an iterative meta-analysis, employing inverse-variance weighted meta-analysis in METAL, separately for UKBB and BBJ cohorts. For the meta-analysis of GWAS results derived from single-ancestry analyses in the UKBB and BBJ (referred to as “Meta”), we incorporated a variable number of EUR bins from UKBB, ranging from 8 to 64 with an increment of 8. Subsequently, we systematically integrated additional EAS bins from BBJ.

##### PRS construction methods

We used different methods to construct PRS in the target populations, specifically UKBB-EAS and UKBB-EUR. In accordance with *Simulations*, we also explored the impact of LD reference panels on PRS performance by utilizing multiple panels from 1KG, while taking into account the ancestry composition of discovery GWAS for P+T. Additionally, we implemented PRS-CS^39^, a Bayesian regression framework that integrates a continuous shrinkage prior to infer the posterior mean effects of SNPs. To ensure computational efficiency, we employed the auto model in the PRS-CS framework, which automatically estimates the hyper-parameter *phi* (the proportion of SNPs with non-zero effects) based on the input GWAS (see **Supplementary Note 8**). For both UKBB and Meta, we used 1KG-EUR as the LD reference panel, while for BBJ, we utilized 1KG-EAS reference panel.

To further explore the performance of PRS incorporating GWAS from multiple ancestries, we constructed a weighted PRS by linearly combining PRS derived from single-ancestry GWAS^34^. Specifically, the weighted PRS was calculated as **PRS_weighted_** = *w*_1_* PRS_EUR_GWAS_ + *w*_2_ * PRS_Minor_GWAS_, where *w*_1_ and *w*_2_ were weights attached to individual PRS. Furthermore, we used a more sophisticated method, PRS-CSx^8^, to generate ancestry-specific posterior SNP effects using multiple GWAS summary statistics. PRS-CSx, an extension of PRS-CS, can model ancestry-specific allele frequencies and LD patterns. Similar to PRS-CS, we used the ancestry-matched LD reference panel from 1KG and performed the auto model implemented in PRS-CSx. We also incorporated the *--meta* flag, which enables inverse-variance weighted meta-analysis in the Gibbs sampler. Consequently, we developed two types of PRS from PRS-CSx, one was based on the meta-analyzed effects (referred to as PRS_multi_) and the other, PRS_weighted_, was dependent on the ancestry-specific posterior SNP effects.

##### PRS performance evaluation

We assessed the predictive accuracy of PRS by measuring the incremental *R*^2^ using linear regression, where we accounted for the influence of covariates. Two models were compared: 1) *H*_0_: *Phenotype* ~ *covariates*, representing the baseline model, and 2) *H*_1_: *Phenotype* ~ *PRS + covariates*, incorporating PRS as the full model. The incremental *R*^2^ was utilized to quantify the improvement in model accuracy resulting from the inclusion of PRS, thus providing a measure of the specific contribution made by PRS to the predictive power of the model. We computed the corresponding 95% confidence intervals (CIs) through bootstrap. To maximize the predictive accuracy of P+T and PRS_weighted_, we employed an optimization strategy to identify the optimal *p*-value thresholds for P+T and the weights (*w*_1_ and *w*_2_) assigned to various PRS components for PRS_weighted_. This optimization process entailed a random partitioning of the target population into two equally sized subsets, namely the validation dataset and the test dataset. The hyperparameter was identified in the validation dataset, and subsequently, the accuracy of the model was assessed using the test dataset. We replicated the process 100 times and calculated the standard error of predictive accuracy across 100 replicates. This approach allowed us to maximize the performance of P+T and PRS_weighted_ by iteratively refining the *p*-value thresholds and weight parameters, thereby enhancing their predictive capabilities.

#### Measures of genetic architecture using summary-data-based BayesS (SBayesS)^29^

To better understand the impact of trait genetic architecture on PRS predictive performance, we evaluated three parameters including the polygenicity (proportion of SNPs with nonzero effects), SNP-based heritability and *S* (the relationship between MAF and effect sizes) for 17 studied phenotypes (**Table S3)**. These parameters were estimated using SBayesS implemented in the GCTB software (https://cnsgenomics.com/software/gctb/). For the analysis, we employed meta-analyzed GWAS data obtained from the comprehensive UKBB and BBJ datasets. Specifically, the number of bins included in the GWAS was equal to the total number of bins associated with the respective phenotype (**Table S3)**. We used the LD reference panel provided by GCTB for UKBB GWAS. We constructed a shrunk LD matrix using 50,000 unrelated individuals from BBJ as the LD reference panel for BBJ GWAS. We used 4 chains for the Markov Chain Monte Carlo process, which calculated the Gelman-Rubin convergence diagnostic (also known as potential scale reduction factor) for these three parameters. We performed the analyses using other default settings for SBayesS. Given the potential convergence issues associated with Bayesian models, we deemed a threshold value of less than 1.2 for the Gelman-Rubin convergence diagnostic as indicative of good convergence for the estimated parameters.

### UK Biobank recent admixture ancestry analysis

To investigate one explanation for poor transferability of PRS across populations – genetic divergence between the discovery and target cohorts – we further explored whether PRS constructed from ancestry-specific summary statistics generated with local ancestry-informed GWAS in admixed populations improves predictive performance in underrepresented populations. Specifically, we used the Tractor method^19^, accounting for both local ancestry and risk allele information, to run GWAS in two-way admixed AFR-EUR individuals from the UKBB (N = 4,576). The average AFR proportion was 62.9%. We used 4,022 unrelated relatively homogeneous AFR individuals, which are independent from the admixed individuals, as the target cohort.

We followed the same criteria for QC and individual selection as described in Atkinson et al.^19^. For sample QC, we excluded individuals that had <95% call rate, withdrew from the study, had closer than 2nd degree relatives present in the sample, or that had sex chromosome aneuploidies. For variant QC we restricted to biallelic SNPs with >90% call rate, HWE *p*-value > 10^-6^, and MAF of at least 0.5%. We selected two-way admixed AFR-EUR individuals from the UKBB by first using the PC loadings from the reference dataset described previously for ancestry inference (1KG + HGDP) to project UKBB individuals into the same PC space. We applied the same random forest ancestry classifier described previously to the projected UK Biobank PCA data and assigned AFR ancestry if the probability was >50%. We restricted to only two-way admixed AFR-EUR ancestry individuals by selecting those individuals assigned the ‘AFR’ population label, then filtering to those with at least 12.5% European ancestry, at least 10% African ancestry, and who did not deviate more than 1 standard deviation from the AFR-EUR cline based on their PC loadings. This process resulted in 4,576 individuals.

We ran local ancestry deconvolution on this set of admixed individuals using RFmix v2^18^ with 1 EM iteration and a window size of 0.2 cM with the HapMap combined recombination map^57^ to inform switch locations. The-n 5 flag (terminal node size for random forest trees) was included to account for an unequal number of reference individuals per reference population. We used the -- reanalyze-reference flag, which recalculates admixture in the reference samples for improved ability to distinguish ancestries. As a reference panel, we used continental AFR and EUR individuals from the 1KG.

Subsequently, we performed GWAS for the 17 quantitative traits utilizing the Tractor method on the 4,576 individuals with mixed AFR-EUR ancestry from the UKBB. This analysis yielded the generation of ancestry-specific summary statistics for the AFR (AFR_Tractor_) and EUR (EUR_Tractor_) ancestry components. To evaluate the performance of PRS in the UKBB-AFR, we developed PRS using Tractor GWAS. Furthermore, we compared these local-ancestry informed PRS with those derived from GWAS conducted using standard methodologies. Specifically, we constructed PRS using GWAS performed on the same set of admixed individuals utilizing the simple linear regression model (ADM_Standard_). Additionally, GWAS summary statistics obtained from UKBB (EUR_standard_, N = 320,000) from the previous section were utilized, and a meta-analysis was conducted to combine the AFR_Tractor_ with EUR_standard_ (Meta_standard_, N = 324,576). We constructed PRS based on HapMap3 SNPs, as previous studies have shown comparable performance between using reliable HapMap3 SNPs exclusively and the use of genome-wide SNPs^7, 58^. Additionally, we constructed weighted PRS by incorporating GWAS of AFR_Tractor_ and EUR_Standard_, for P+T, PRS-CS and PRS-CSx, respectively. Considering the ancestry composition of the discovery GWAS, we used different sets of reference panels for each respective GWAS. Specifically, we used 1KG-EUR as the LD reference panel for EUR_Tractor,_ EUR_standard_ and Meta_standard_, while using 1KG-AFR for AFR_Tractor_. We used an in-sample LD panel for ADM_Standard_. We calculated the predictive accuracy in the UKBB-AFR using incremental *R*^2^ as described above. We repeated the process 100 times and reported the standard error of predictive accuracy across 100 estimates.

## Excel Table Title and Legends

Table S1. The comparison of using different LD reference panels across various simulation scenarios. Related to Figure S2, Figure 2, Figure S3 and Figure S4.

Table S2. Impact of cross-ancestry genetic correlation on predictive performance. Related to Figure S5.

Table S5. Predictive accuracy for P+T and PRS-CS across phenotypes using single-ancestry discovery GWAS from UKBB and BBJ. Related to Figure S6 and Figure S11.

Table S6. Impact of LD reference panel on P+T performance using multi-ancestry GWAS for 17 traits. Related to Figure S7, Figure S8 and Figure S11.

Table S7. Accuracy differences between using PRS derived from multi-ancestry GWAS and using PRS derived from EUR GWAS. Related to Figure S9 and Figure S10.

Table S8. Accuracy differences between PRS derived from multi-ancestry GWAS and using PRS from weighted linear combination. Related to Figure 5 and Figure S12.

Table S9. Predictive accuracy in the UKBB using PRS-CSx for 17 traits. Related to Figure 5 and Figure S12.

Table S10. Predictive accuracy in the UKBB-AFR using various discovery GWAS. Related to Figure 6.

## Notes

### Summary of Updates

Manuscript and figures

