## Supplementary Notes, Figures and Tables for "Polygenic prediction across populations is influenced by ancestry, genetic architecture, and methodology"

#### 1. Impact of LD reference panels on predictive accuracy of PRS constructed from multi-ancestry GWAS in simulations

We first investigated the impact of different LD reference panels on the predictive accuracy of PRS when the ancestry composition of the multi-ancestry GWAS varied. We utilized three sets of LD reference panels, including two single-ancestry datasets (N=10,000) matching the ancestry composition of each contributing population to the discovery GWAS, and one combined dataset (N=10,000) with individuals proportional to the ancestry composition of the discovery GWAS.

Our findings showed that the impact of the LD reference panel was more subtle for more polygenic traits compared to less polygenic ones (**Figure S2** and **Table S1**). Using different reference panels, the median predictive accuracies in more polygenic scenario ($M_c$ = 1000 and $h^2$ = 0.03) were ~0.014 in AFR and ~0.017 in EAS, respectively. While, such accuracies ranged from 0.029 to 0.033 in AFR and from 0.038 to 0.042 in EAS for the less polygenic scenario with $M_c$ = 100 and $h^2$ = 0.05. When using single-ancestry LD reference panels, we found that using the one matching the majority ancestry in the discovery GWAS yielded better predictive performance. For example, when there were more EUR samples, the median accuracy in AFR across simulations was 0.023 using EUR LD reference compared to 0.016 using AFR LD reference. Furthermore, we found that such single-ancestry LD reference panels generally provided comparable predictive accuracy to the proportional combined ancestry panel (referred to as “Combined”, **Figure S2**), especially when the ancestry composition was increasingly disproportionate. In particular, the Combined LD panel did not yield significantly better PRS accuracy compared to the optimal single-ancestry LD panel, and minor differences were smallest for the most polygenic scenarios. Our simulation setup allowed for the proportion of understudied populations to exceed 50%, although this was not always the case in current multi-ancestry GWAS. Therefore, we reported our results based on estimates using the Combined LD reference panel to avoid arbitrariness when ancestry proportions for multi-ancestry GWAS are similar.

#### 2. Predictive performance of PRS derived from multi-ancestry GWAS in simulations

We observed consistent upward trends of predictive accuracy in the understudied target populations with increasing target-ancestry matched samples included in discovery GWAS (**Figure S2**). The extent of the improvement varied depending on the genetic architecture of the trait, with more polygenic traits exhibiting greater benefits. In comparison to including only one bin from the target population, we observed the largest accuracy improvement in AFR of 0.006, 0.006 and 0.011 for $M_c$ of 100, 500, and 1000, respectively, and of $h^2$ of 0.03.

Furthermore, we found that for less polygenic traits with a larger per-variant explained variance, the accuracy plateaued sooner when fewer bins from minority populations were included compared to more polygenic traits with lower variance explained for each variant. As shown in

**Figure S2**, the highest accuracy in AFR was achieved with the inclusion of 7 AFR bins in the discovery GWAS for the least polygenic scenario ( $M_c$ = 100 and $h^2$ = 0.05). For the most polygenic scenario ($M_c$ = 1000 and $h^2$ = 0.03), the accuracy continued to improve as more AFR bins were included.

#### 3. Relative accuracy using $PRS_{multi}$ compared to using $PRS_{EUR\_GWAS}$

Specifically, the relative accuracy here was calculated as the difference in PRS $R^2$ between the PRS derived from multi-ancestry GWAS and EUR GWAS divided by the PRS $R^2$ in EUR ancestry from the EUR GWAS, i.e. $RA = \frac{R^2_{target}\ using\ PRS_{multi} - R^2_{target}\ using\ PRS_{EUR\_GWAS}}{R^2_{EUR}\ using\ PRS_{EUR\_GWAS}}$. Therefore, the trend of RA was consistent with the accuracy improvement of $PRS_{multi}$.

#### 4. PRS accuracy improvement will diminish with multi-ancestry GWAS compared to using Minor GWAS only

In comparison to PRS derived from Minor GWAS alone ($PRS_{Minor\_GWAS}$), we found that the accuracy improvement of $PRS_{multi}$ gradually diminished as the sample size of Minor GWAS increased (**Figure S4 and Table S1**). We showed that in general little to no improvement was achieved by $PRS_{multi}$ when the understudied target populations accounted for more than half sample sizes of the multi-ancestry GWAS for more polygenic traits. However, for the less polygenic traits, a much smaller Minor GWAS outperformed large-scale multi-ancestry GWAS. For instance, in the least polygenic scenario ( $M_c$ = 100 and $h^2$ = 0.05), no improvement of $PRS_{multi}$ was observed in EAS with the inclusion of more than 4 EAS bins, and in AFR with more than 2 AFR bins included (**Figure S4)**. On the other hand, we observed slight improvements of 0.002 in the target populations when more than 16 target ancestry-matched bins were included in the most polygenic scenario ( $M_c$ = 1000 and $h^2$ = 0.03). Interestingly, we observed consistent accuracy improvements for target populations of EUR and the ancestry not included in the multi-ancestry GWAS, although such improvement decreased with larger numbers of bins from minority populations. This could be due to the multi-ancestry GWAS being more genetically similar to those populations compared to the Minor GWAS.

#### 5. Impact of cross-ancestry genetic correlation on PRS predictive accuracy using multi-ancestry GWAS in simulations

By relaxing the assumption of perfect correlation between effect sizes of causal variants across populations, we expanded two simulation scenarios, scenario 1 with $M_C$ = 100 and $h^2$ =0.05, and scenario 2 with $M_C$ = 1000 and $h^2$ =0.03, by simulating different values of pairwise cross-ancestry genetic correlation ($r_g$ = 0.6 and 0.8). We subsequently investigated the impact of $r_g$ on the accuracy of $PRS_{multi}$ compared to $PRS_{single}$ (**STAR Method**).

Consistent with our previous findings when $r_g$ was 1, we observed consistent improvements in accuracy for the more polygenic scenario 2 in the target population when including more individuals from that specific ancestry in multi-ancestry GWAS, as compared to the accuracy of $PRS_{EUR\_GWAS}$ (**Table S2 and Figure S5-A,B**). Moreover, for the less polygenic scenario 1, achieving near-saturated accuracy improvement using $PRS_{multi}$ required a larger number of individuals from the understudied target population, especially for traits with lower $r_g$. For instance, with the inclusions of more than 34, 24 and 15 EAS bins, the accuracy improvements in EAS reached approximately 0.019, 0.010 and 0.008 for $r_g$ values of 0.6, 0.8 and 1, respectively.

Furthermore, when comparing the accuracy of $PRS_{multi}$ to $PRS_{Minor\_GWAS}$, we observed little to no improvements, particularly when the sample size of the Minor GWAS was larger and the $r_g$ was lower (**Figure S5-C,D)**. This trend was more pronounced in AFR, likely attributed to their greater genetic diversity and larger genetic divergence from EUR. These findings underscore the necessity of increasing the diversity of genomic studies and enhancing sample sizes of underrepresented populations.

#### 6. PRS accuracy using smaller target ancestry-matched GWAS versus larger-scale EUR GWAS may be comparable depending on methodology and trait-specific genetic architecture

We constructed PRS using P+T and PRS-CS for different phenotypes in the target populations using single-ancestry GWAS from UKBB and BBJ, respectively. The GWAS sample size varied depending on the number of total bins for that phenotype, with each bin comprising 5,000 individuals randomly sampled from the respective dataset (**Table S3**).

Overall, there was a clear increasing trend in the target populations between PRS accuracy and a larger discovery GWAS (**Figure S6** and **Table S5**). However, such patterns differed by ancestry and PRS methods in a trait-specific manner. For example, in the UKBB-EAS the upward trend was not obviously witnessed for basophil, a rare cell type, using BBJ where the predictive accuracy was not significantly different from 0. This observation might be attributed to factors such as smaller GWAS sample sizes, ascertainment bias and low heritability in the BBJ. Moreover, we observed that the more sophisticated method PRS-CS generally significantly outperformed the classic P+T method across traits especially for traits with higher polygenicity and larger sample sizes (one-side Wilcoxon test, $p$-value < 0.05). Specifically, the median accuracy of PRS derived from BBJ in the UKBB-EAS was 0.013 and 0.010 using PRS-CS and P+T, respectively. The corresponding values were 0.046 and 0.032 when the discovery GWAS was UKBB. However, we observed specific cases where P+T outperformed PRS-CS. For MCH and MCV with ancestry-enriched variants exhibiting large effects, using P+T with BBJ yielded higher accuracy compared to PRS-CS. Further, we compared the performance of P+T and PRS-CS using the full BBJ and UKBB, respectively, in the UKBB-EAS population. Despite the substantially smaller sample size in BBJ, the accuracy of PRS using P+T was comparable to that of PRS using full UKBB across various traits (median $R^2$: 0.019 VS 0.038, one-sided Wilcoxon signed-rank test, $p$-value = 0.171). However, using PRS-CS with full UKBB exhibited significantly higher accuracies across traits compared to using PRS-CS with the full BBJ (median $R^2$: 0.051 VS 0.018, one-sided Wilcoxon signed-rank test, $p$-value = 0.001).

Consistent with previous work[1–4], $PRS_{single}$ was generally more transferable (as measured by relative accuracy, the ratio of predictive accuracy between target populations) when the target population was more genetically related to the discovery GWAS (**Figure S7**). Interestingly, we observed that in comparison with predictive accuracy, the relative accuracy of PRS did not show a clear increase with larger sample sizes in UKBB. These results suggest that the PRS transferability issue is unlikely to be addressed by solely relying on larger EUR GWAS.

#### 7. Predicted accuracy in cross-ancestry polygenic prediction using different discovery GWAS.

The expected accuracy of PRS in the UKBB-EAS derived from BBJ is based on the theoretical equation: $R^2 \approx \frac{h_d^2}{h_d^2 + \frac{M_d}{N_d}}$ (1)[5], where $h_d^2$ denotes the SNP-based heritability in the discovery population, $N_d$ is the discovery GWAS sample size and $M_d$ is the number of independent chromosome segments in the discovery population, which we assume to be 50,000[6]. The results are shown in **Figure S14**. When there is imperfect cross-ancestry genetic correlation ($r_g$), we used a generalization of this formula: $R^2 \approx \frac{r_g^2 h_d^2 h_t^2}{h_d^2 + \frac{M_d}{N_d}}$ (2)[7], where $h_t^2$ denotes SNP-based heritability in the target populations. It is important to note that this formula provides an approximation and does not explicitly account for differences in LD structure between ancestries, except for the influence of LD disparities on genetic correlation.

#### 8. Predictive performance of PRS-CS using different models in the UKBB

To alleviate computational burdens, we initially ran PRS-CS using GWAS summary statistics from UKBB with varying numbers of bins (ranging from 8 to 64, with an increment of 8) for 17 traits. We systematically explored the influence of the hyper-parameter (*phi*), representing the proportion of SNPs with non-zero effects, on PRS performance, considering diverse GWAS sample sizes and trait genetic architectures. Specifically, we performed both the grid model with various *phi* parameters ($1\times 10^{-6}$, $1\times 10^{-4}$, 0.01 and 1) and the auto model, which automatically estimates the *phi* parameter based on the input GWAS. We used default settings for all other parameters. Our findings indicated that PRS-CS-auto exhibited comparable predictive accuracy across all traits in the UK Biobank dataset when compared to using the optimal phi parameter in the grid model (**Figure S15**).

### Supplementary Figures

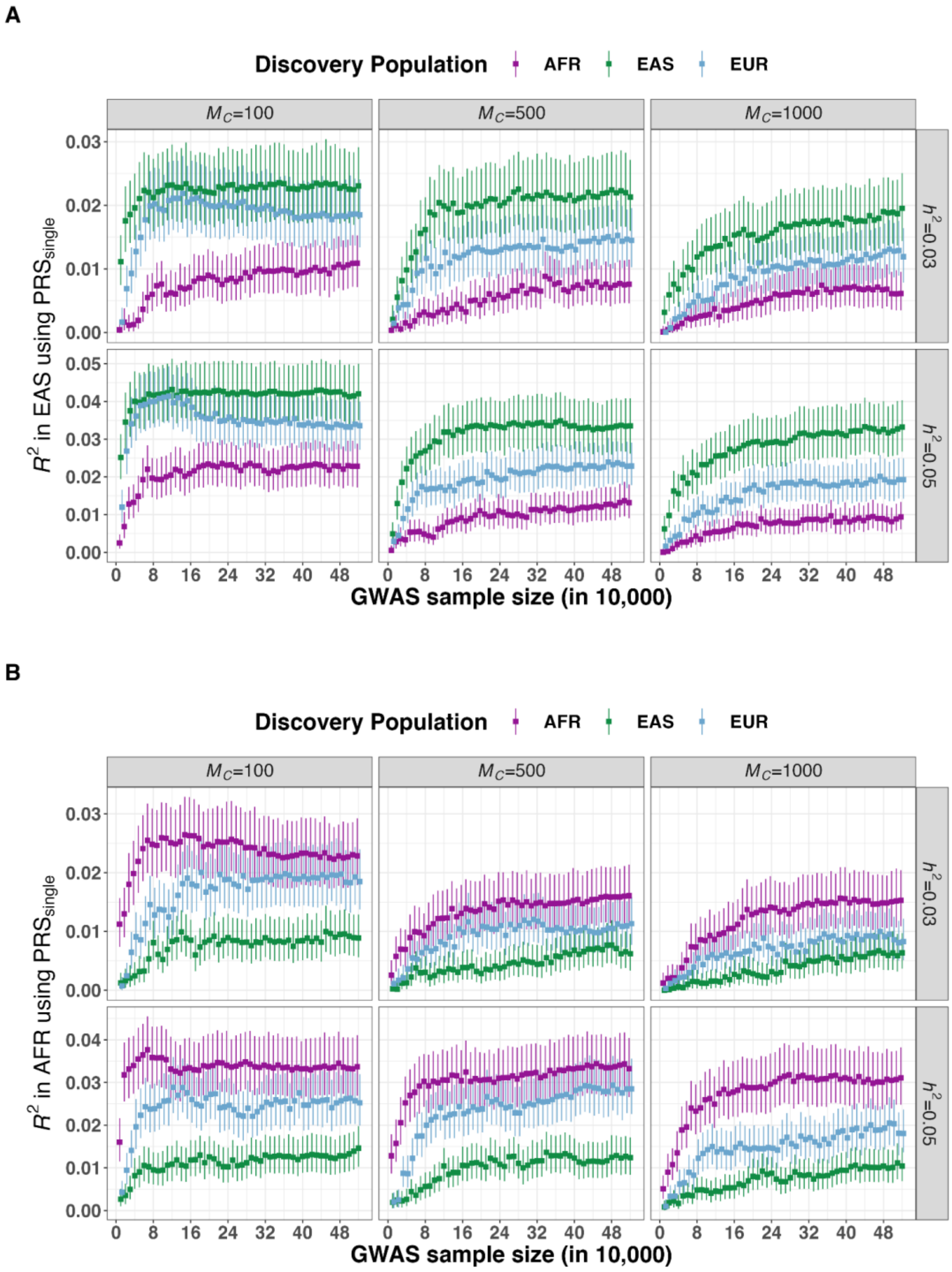

#### Figure S1. Predictive accuracy in the target population using various single-ancestry discovery GWAS in 6 simulated genetic architectures.

We evaluated the accuracy of PRS using P+T in A) East-Asian (EAS) population and B) African (AFR) population using discovery GWAS from AFR, EAS and European (EUR) populations, respectively. $M_c$ indicates the number of causal variants and $h^2$ refers to SNP-based heritability. Full results are shown in **Table S1**. The error bars represent the 95% CIs of the predictive accuracy.

A

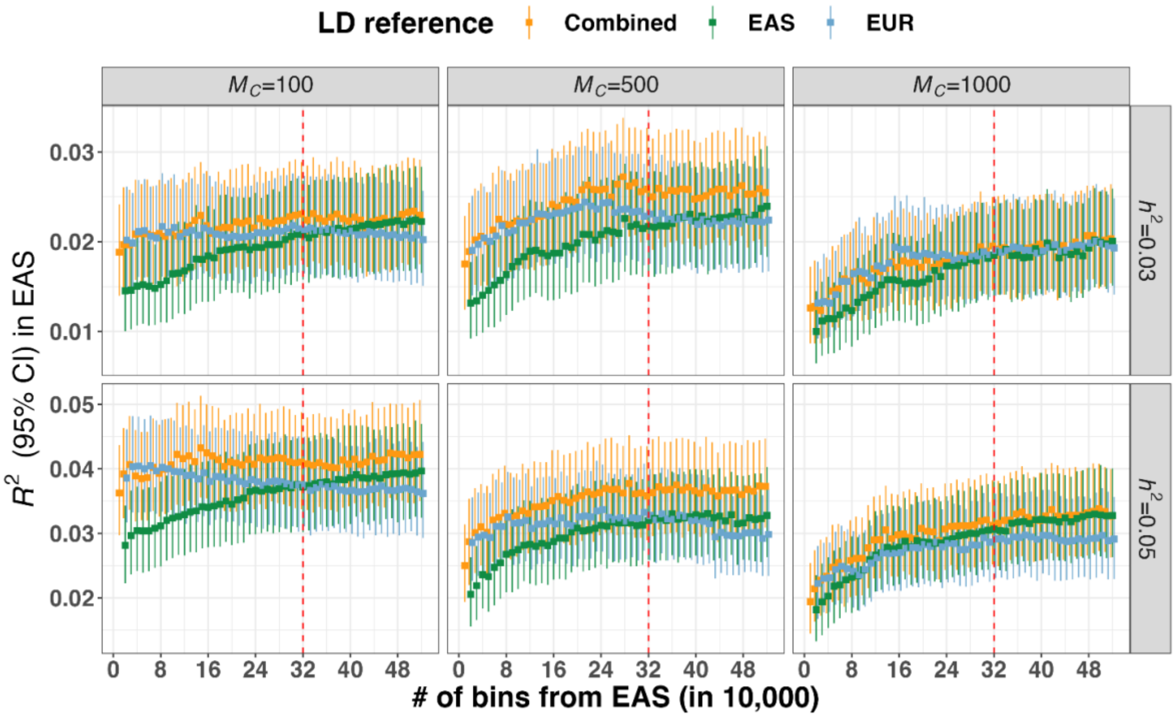
LD reference
Combined
EAS
EUR
$M_C$=100
$M_C$=500
$M_C$=1000
$h^2$=0.03
$h^2$=0.05
$R^2$ (95% CI) in EAS
### of bins from EAS (in 10,000)

B

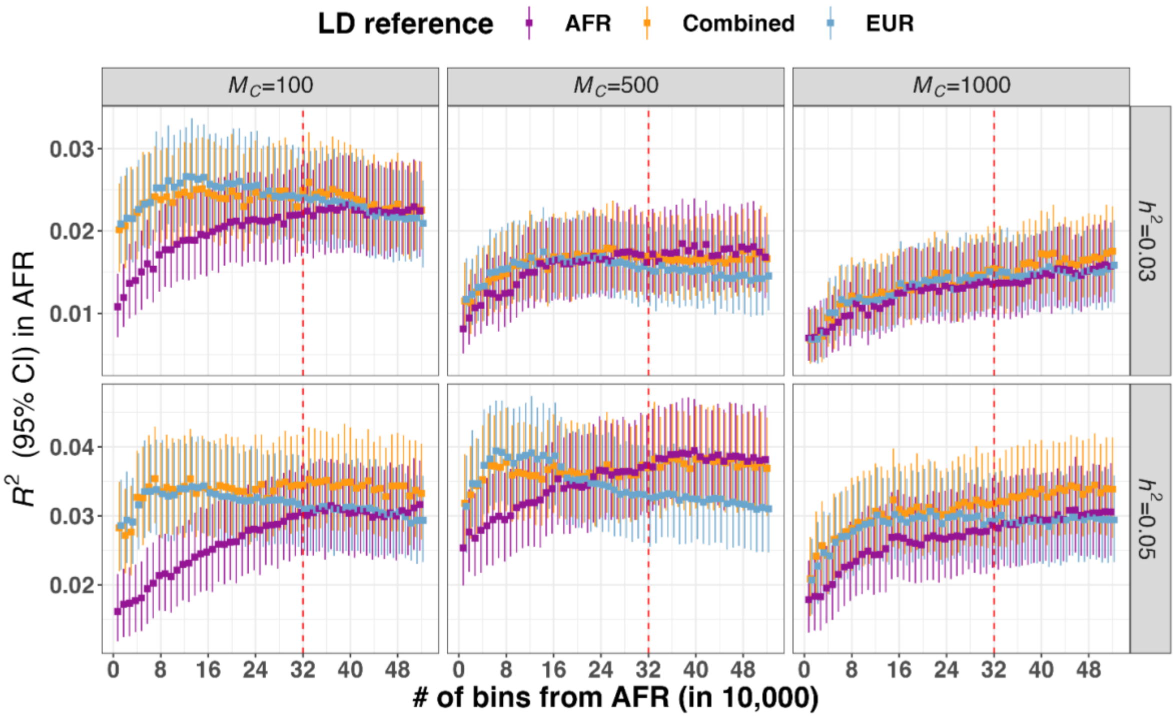
LD reference
AFR
Combined
EUR
$M_C$=100
$M_C$=500
$M_C$=1000
$h^2$=0.03
$h^2$=0.05
$R^2$ (95% CI) in AFR
### of bins from AFR (in 10,000)

#### Figure S2. Impact of LD reference panels on the predictive accuracy using PRS derived from multi-ancestry GWAS.

The multi-ancestry GWAS included populations of European (EUR) and East-Asian (EAS) or African (AFR) ancestry, with the EAS or AFR sample size varying as indicated on the x-axis. Each bin included 10,000 individuals. For illustrative purposes, we present the results where 32 EUR bins were included in the multi-ancestry GWAS. $M_c$ indicates the number of causal variants and $h^2$ refers to SNP-based heritability. A) The predictive accuracy of PRS using P+T with different LD reference panels in EAS when using discovery GWAS which was meta-analyzed on EUR and EAS populations. B) The predictive accuracy of P+T using P+T with different LD reference panels in AFR when using discovery GWAS which was meta-analyzed on EUR and AFR populations. The error bars represented the 95% CI of the predictive accuracy. For each panel, we used three different LD reference panels, where EUR, EAS and AFR referred to using 10,000 individuals from the ancestry-matched target populations and Combined was denoted as using 10,000 individuals with ancestry proportional to discovery GWAS. The red dashed line in each panel indicates the point where the number of bins from EUR and other populations is the same. Full results are shown in **Table S1**.

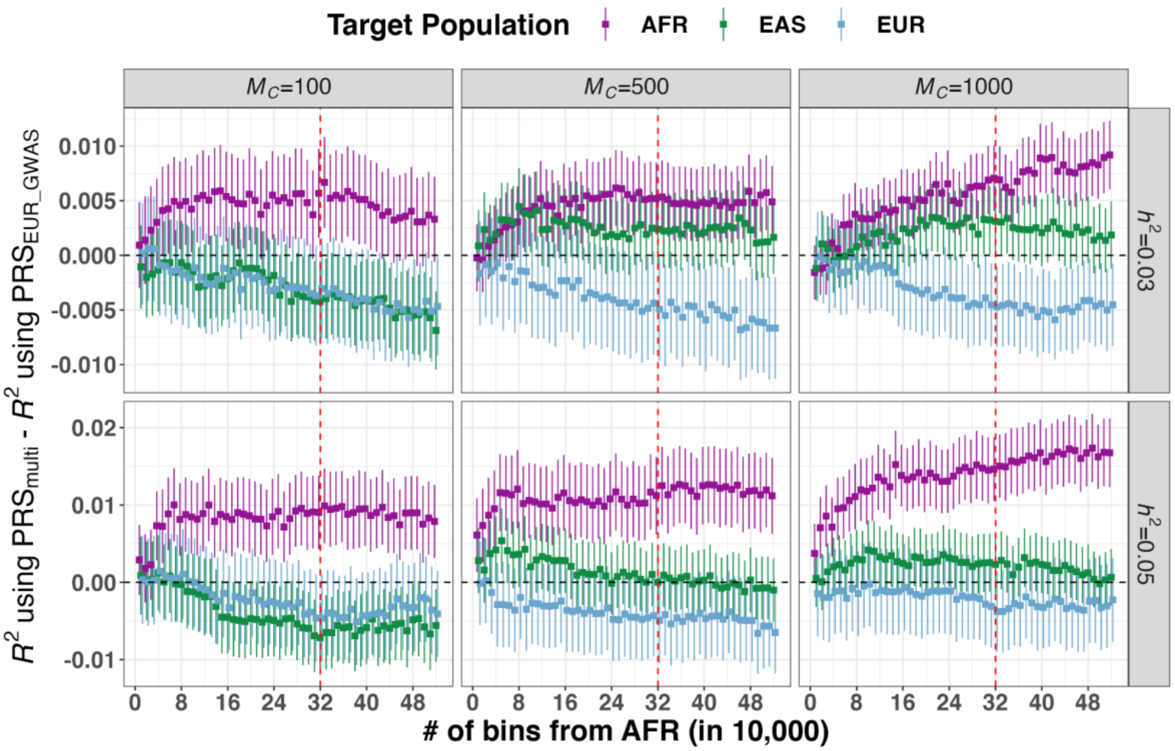

Figure S3. Predictive accuracy improvement of PRS using meta-analyzed multi-ancestry GWAS compared to using large-scale European GWAS in 6 simulated genetic architectures.

The multi-ancestry GWAS included populations of European (EUR) and African (AFR) ancestry, with the AFR sample size varying as indicated on the x-axis. For illustrative purposes, we present the results using 32 EUR bins, each consisting of 10,000 individuals, which were included in both EUR GWAS and multi-ancestry GWAS. PRS was evaluated in AFR, EAS and EUR, respectively. Full results are shown in **Table S1**. $M_c$ indicates the number of causal variants and $h^2$ refers to SNP-based heritability. The red vertical dashed line in each panel indicates the point where the number of bins from EUR and AFR populations is the same. The black horizontal dashed line indicates y=0. The error bars represent the standard errors of predictive accuracy differences using PRS derived from multi-ancestry GWAS ($PRS_{multi}$) and EUR GWAS ($PRS_{EUR\_GWAS}$), respectively.

A

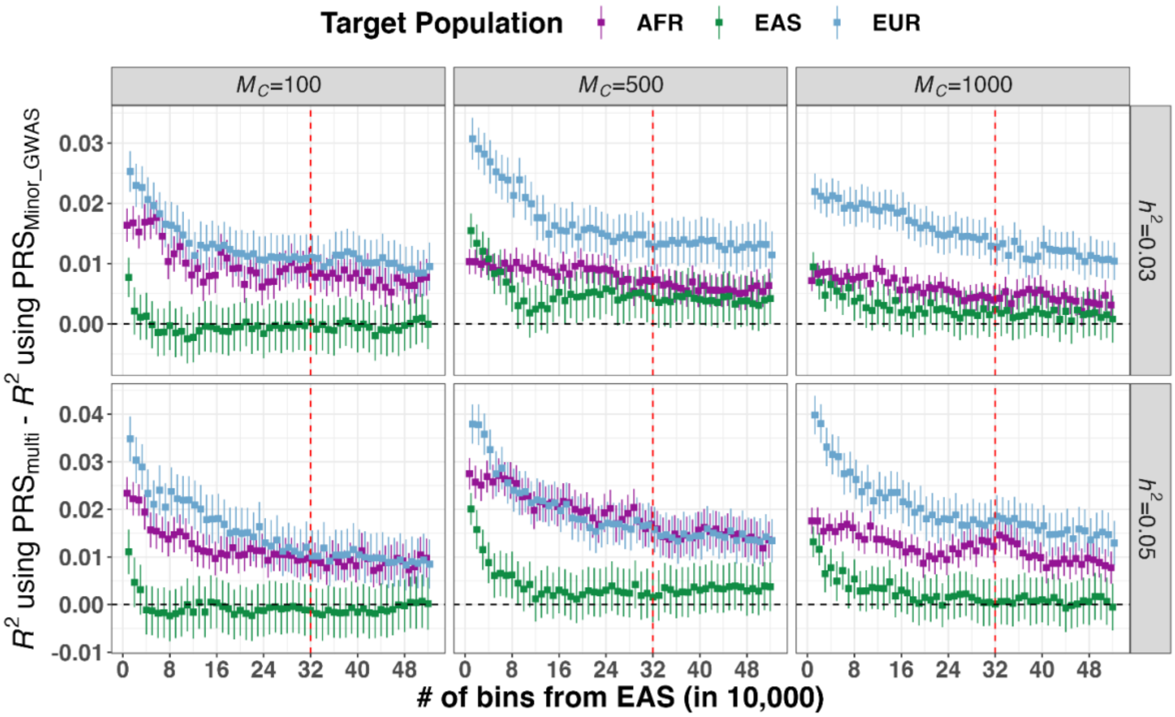

Target Population AFR EAS EUR
$M_C$=100
$M_C$=500
$M_C$=1000
$h^2$=0.03
$h^2$=0.05
$R^2$ using PRS$_{multi}$ - $R^2$ using PRS$_{Minor_GWAS}$
### of bins from EAS (in 10,000)

B

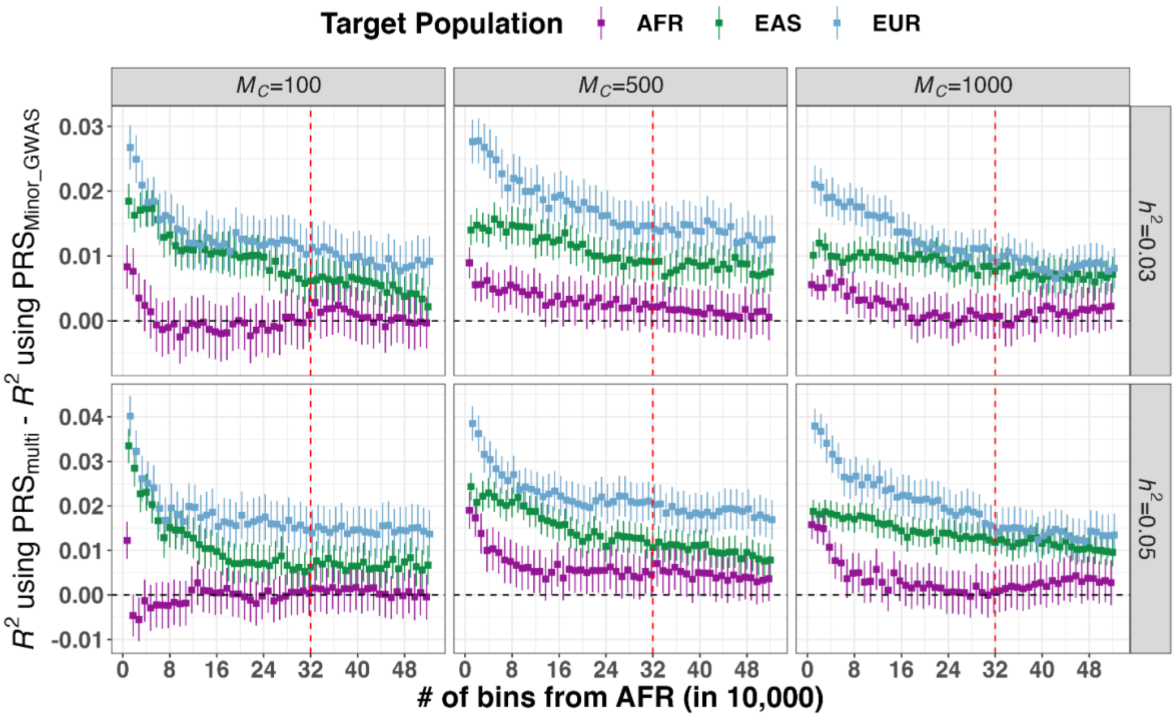

Target Population AFR EAS EUR
$M_C$=100
$M_C$=500
$M_C$=1000
$h^2$=0.03
$h^2$=0.05
$R^2$ using PRS$_{multi}$ - $R^2$ using PRS$_{Minor_GWAS}$
### of bins from AFR (in 10,000)

#### Figure S4. Predictive accuracy improvement of PRS using meta-analyzed multi-ancestry GWAS compared to using minority population-based GWAS in 6 simulated genetic architectures.

The multi-ancestry GWAS included populations of European (EUR) and East-Asian (EAS) or African (AFR) ancestry, with the EAS (A) or AFR (B) sample size varying as indicated on the x-axis. Each bin included 10,000 individuals. For illustrative purposes, we present the results where 32 EUR bins were included in the multi-ancestry GWAS. PRS was evaluated in AFR, EAS and EUR, respectively. Full results are shown in **Table S1**. $M_c$ indicates the number of causal variants and $h^2$ refers to SNP-based heritability. The red vertical dashed line in each panel indicates the point where the number of bins from EUR and minority populations is the same. The black horizontal dashed line indicates y=0. The error bars represent the standard errors of the predictive accuracy differences using PRS derived from multi-ancestry GWAS ($PRS_{multi}$) and minority population-based GWAS ($PRS_{Minor\_GWAS}$), respectively.

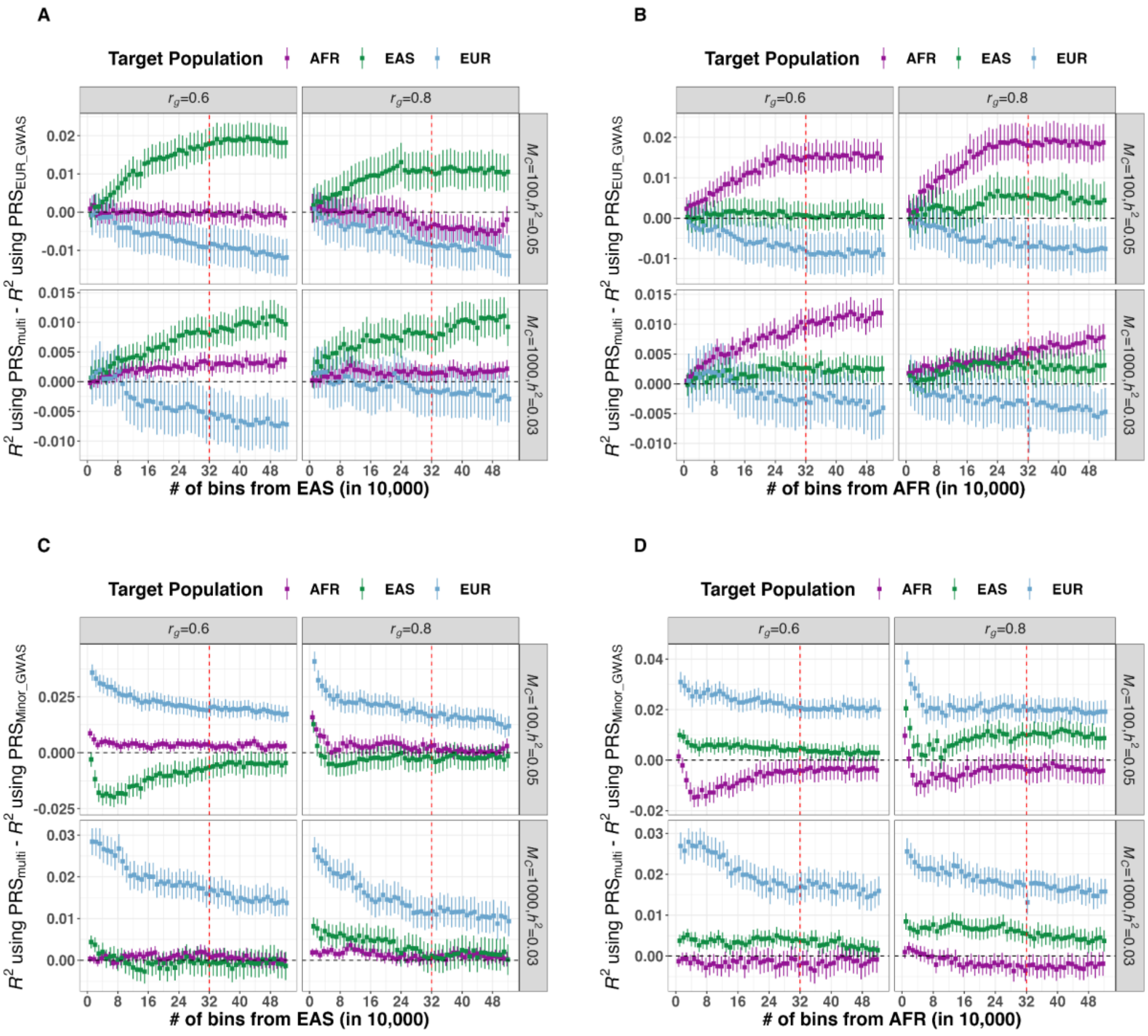

Figure S5. Predictive accuracy improvement of PRS using meta-analyzed multi-ancestry GWAS compared to using single-ancestry GWAS in 2 simulated genetic architectures with varying cross-ancestry genetic correction.

The multi-ancestry GWAS included populations of European (EUR) and East-Asian (EAS) or African (AFR) ancestry, with the EAS (A,C) or AFR (B,D) sample size varying as indicated on the x-axis. Each bin included 10,000 individuals. For illustrative purposes, we present the results where 32 EUR bins were included in both EUR GWAS and multi-ancestry GWAS. PRS was evaluated in AFR, EAS and EUR, respectively. Full results are shown in **Table S2**. $M_c$ indicates the number of causal variants and $h^2$ refers to SNP-based heritability. The red vertical dashed line in each panel indicates the point where the number of bins from EUR and minority populations is the same. The black horizontal dashed line indicates y=0. In A) and B), the error bars represent the standard errors of the predictive accuracy differences using PRS derived from multi-ancestry

GWAS ($PRS_{multi}$) and EUR GWAS ($PRS_{EUR\_GWAS}$), respectively. In C) and D), the error bars represent the standard errors of the predictive accuracy differences using $PRS_{multi}$ and minority population-based GWAS ($PRS_{Minor\_GWAS}$), respectively.

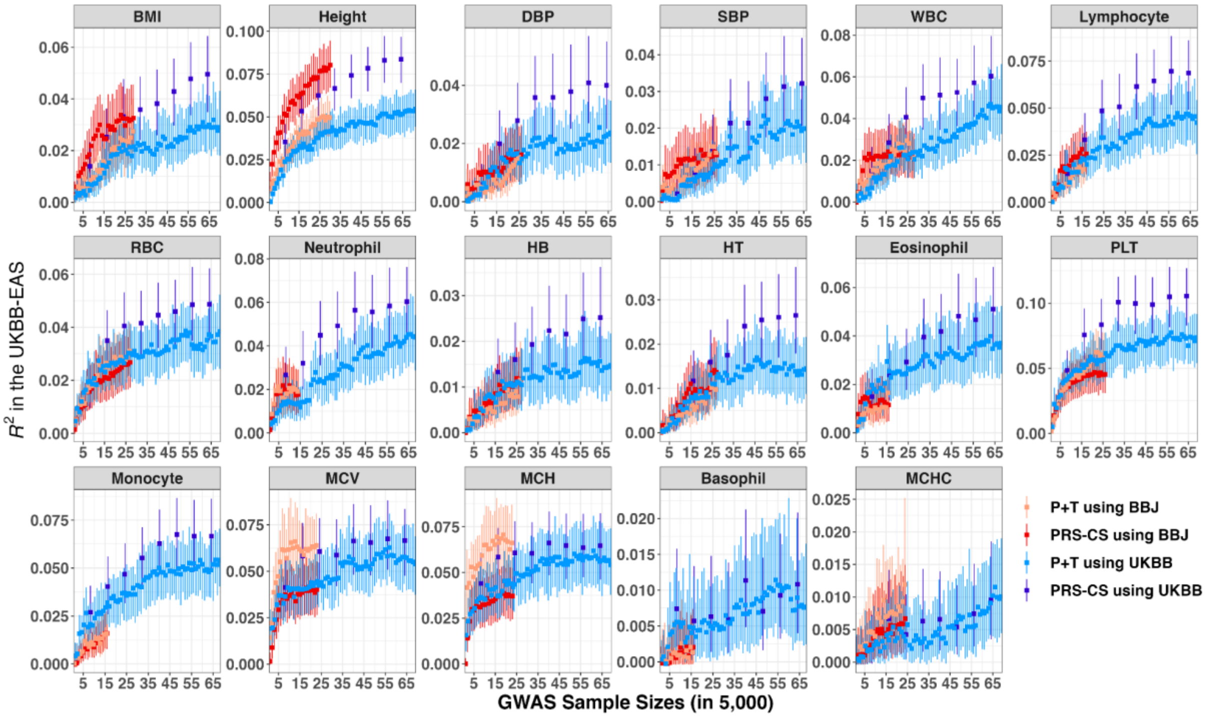

#### Figure S6. Predictive performance in the UK Biobank East-Asian population (UKBB-EAS) using P+T and PRS-CS.

We used GWAS from both Biobank Japan (BBJ) and UK Biobank (UKBB) to construct PRS. We reported the predictive accuracy in the UKBB-EAS using the auto model for PRS-CS and optimal *p*-value for P+T (see **STAR Methods**). The phenotypes were ranked by polygenicity estimated using UKBB (**Figure 3**). The error bars represent the 95% confidence intervals.

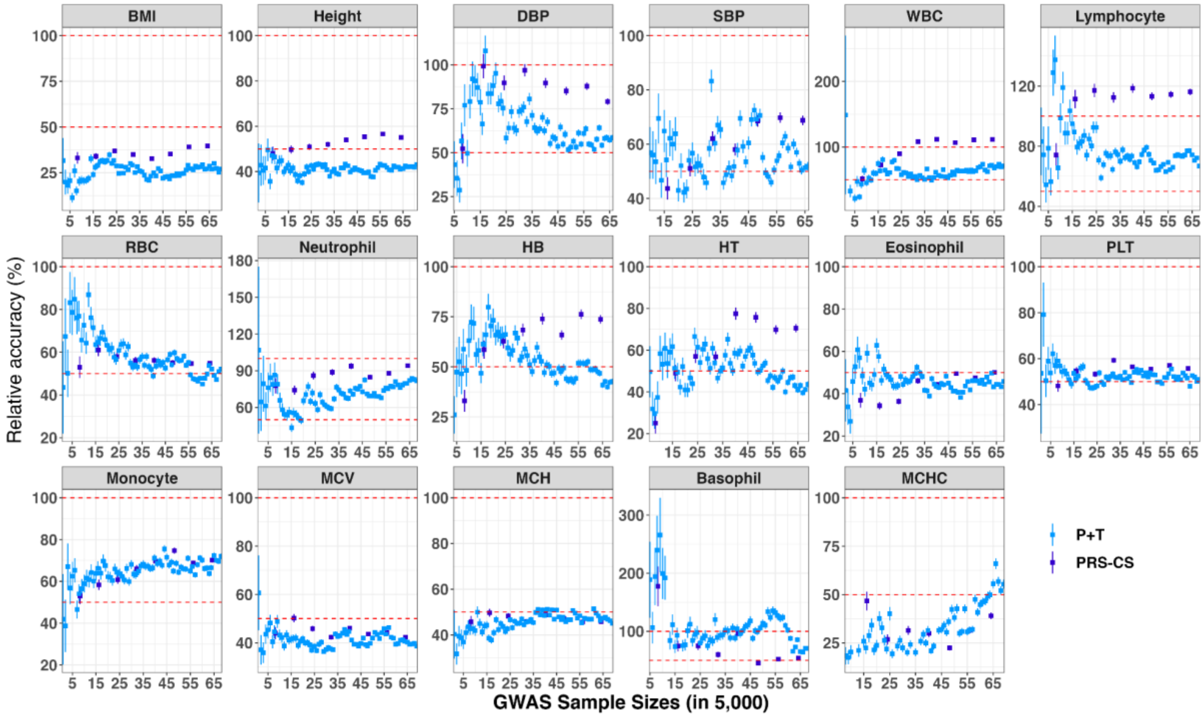

### Figure S7. Relative accuracy using different discovery GWAS with varying sample sizes.

We constructed PRS using both P+T and PRS-CS when derived from UK Biobank (UKBB). We estimated PRS accuracy in both UKBB-EAS and UKBB-EUR. We calculated the relative accuracy as the ratio of accuracy between UKBB-EAS and UKBB-EUR. The error bar represents the standard error of the relative accuracy.

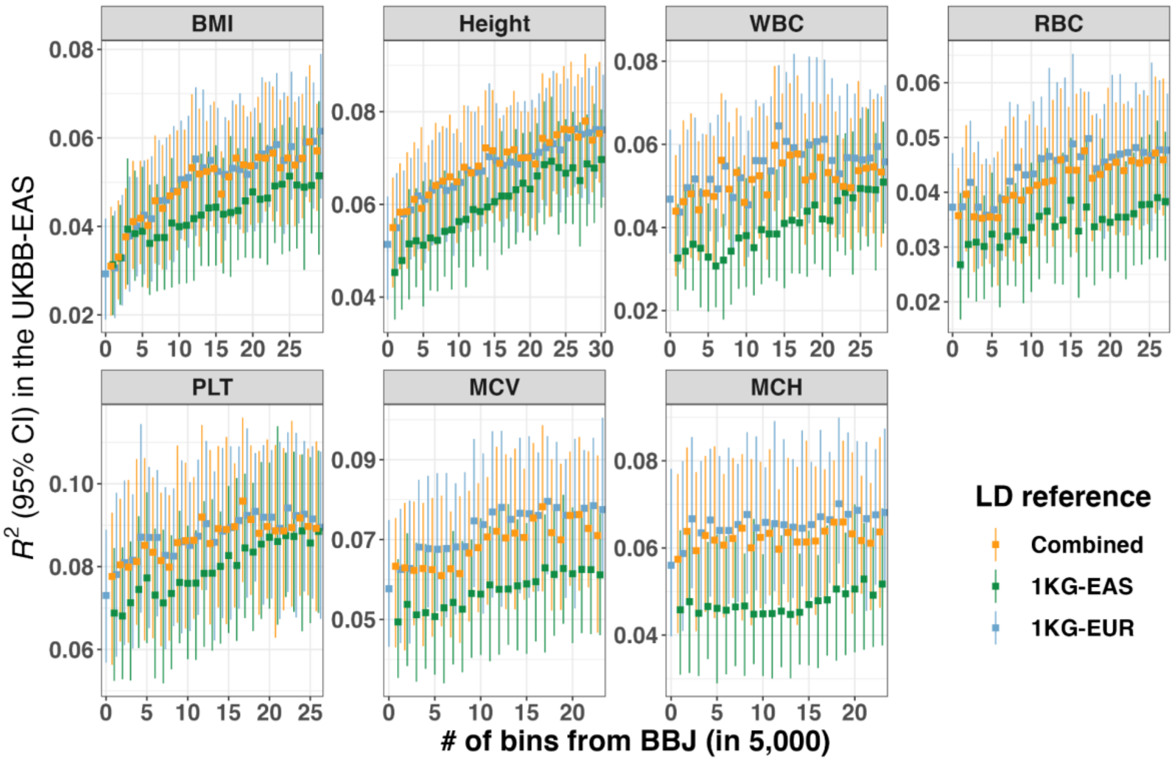

Figure S8. Impact of LD reference on predictive accuracy using P+T for multi-ancestry GWAS across phenotypes.

We used three different LD reference panels, including unrelated European and East-Asian from 1000 Genome Project (1KG-EUR and 1KG-EAS) and combined EUR and EAS (N = 500) from 1KG proportional to the ancestry composition of discovery GWAS, when using P+T for multi-ancestry GWAS. PRS was evaluated in the UKBB-EAS. We showed the results for 7 traits with SNP-based heritability > 0.1 in both Biobank Japan (BBJ) and UK Biobank (UKBB), while they were ranked by polygenicity estimates using UKBB (**Figure 3**). We used 64 bins (N = 320,000) from UKBB as an example. The x-axis is the number of bins from BBJ. The full results are shown in **Table S6**.

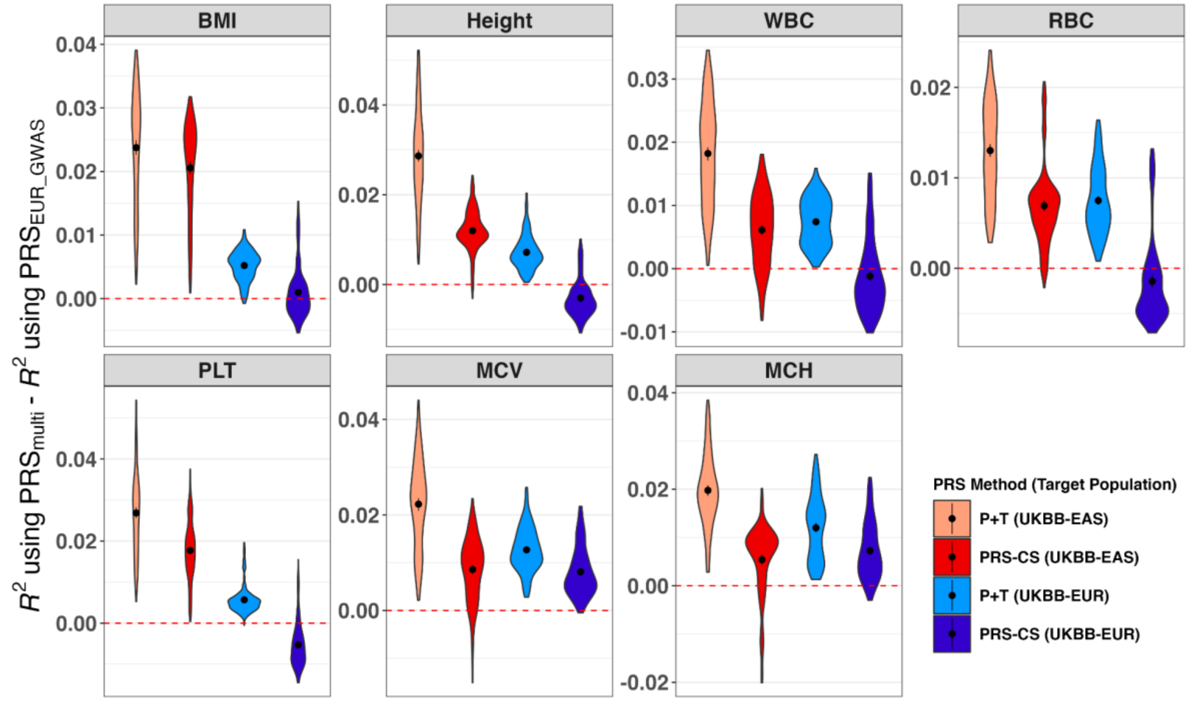

Figure S9. Accuracy improvement of PRS derived from multi-ancestry GWAS relative to from EUR GWAS.

We constructed PRS using P+T and PRS-CS and evaluated them in the UKBB-EAS and UKBB-EUR. The y-axis is the accuracy difference of PRS between using multi-ancestry GWAS ($PRS_{multi}$) and using EUR GWAS ($PRS_{EUR\_GWAS}$) when the number of EUR bins is the same. The error bars indicate the standard error of mean accuracy improvement. The red dashed line is y=0. We showed the results for 7 traits with SNP-based heritability > 0.1 in both Biobank Japan (BBJ) and UK Biobank (UKBB), while they were ranked by polygenicity estimates using UKBB (**Figure 3**). Full results are shown in **Table S7**.

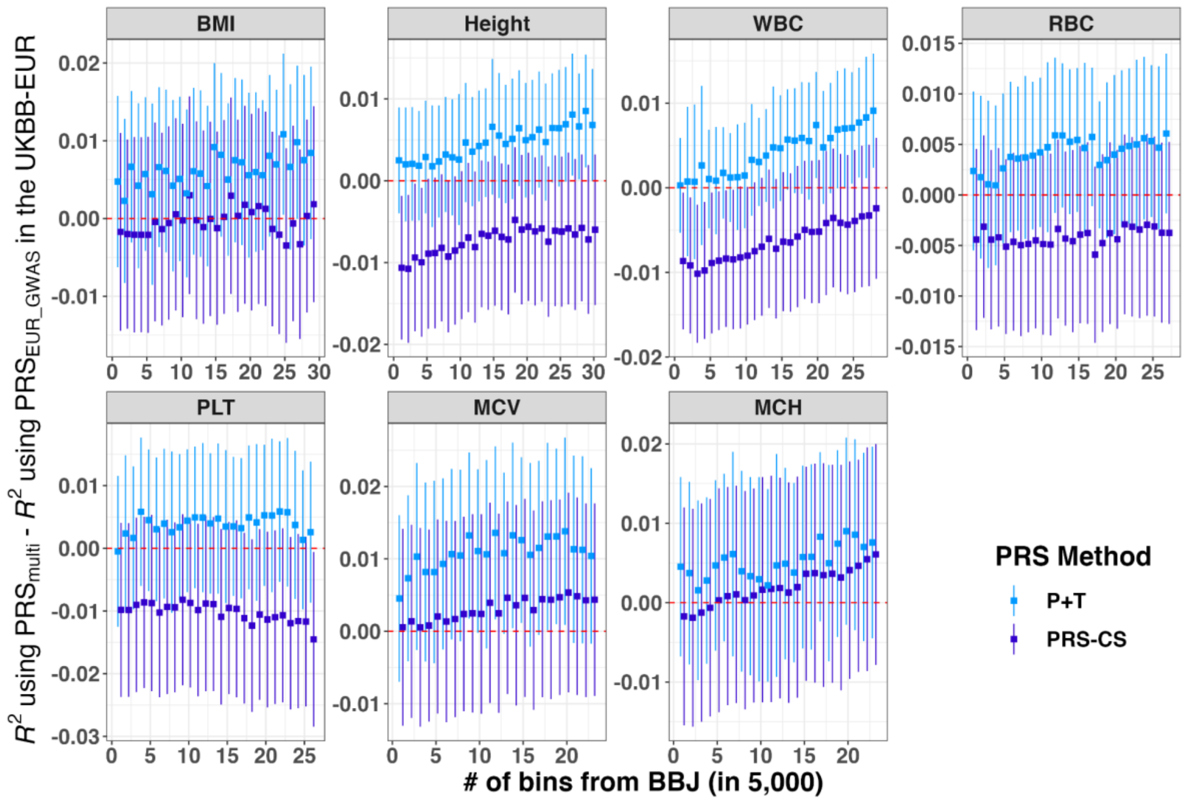

### Figure S10. Accuracy improvement of PRS in the UKBB-EUR using multi-ancestry GWAS relative to using EUR GWAS for P+T and PRS-CS.

We constructed PRS using P+T and PRS-CS and evaluated them in the UKBB-EUR. The y-axis is the accuracy difference of PRS between using multi-ancestry GWAS ($PRS_{multi}$) and using EUR GWAS ($PRS_{EUR\_GWAS}$) when the number of bins from EUR GWAS is 64. The x-axis is the number of bins from BBJ included in the multi-ancestry GWAS. The error bars indicate the standard error of mean accuracy improvement. The red dashed line is y=0. We showed the results for 7 traits with SNP-based heritability > 0.1 in both Biobank Japan (BBJ) and UK Biobank (UKBB), while they were ranked by polygenicity estimates using UKBB (**Figure 3**). Full results are shown in **Table S7**.

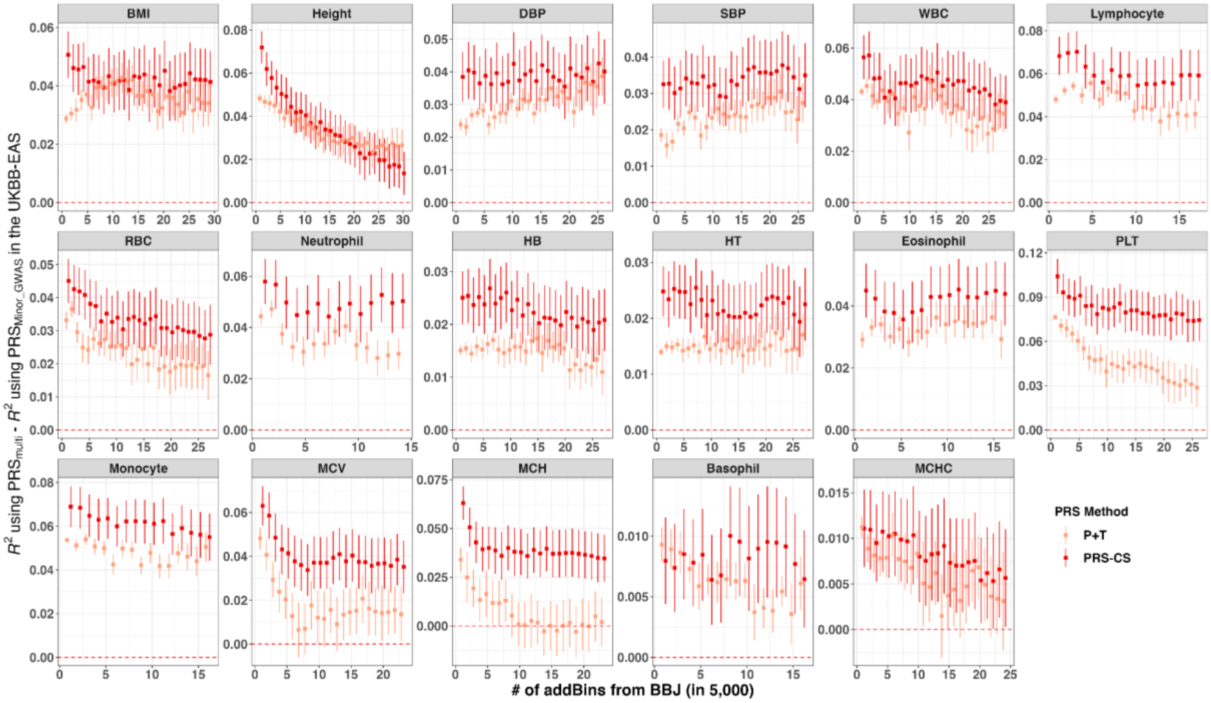

#### Figure S11. Accuracy improvement of PRS in the UKBB-EAS using multi-ancestry GWAS relative to using Minor GWAS for P+T and PRS-CS.

We constructed PRS using P+T and PRS-CS and evaluated them in the UKBB-EAS. The y-axis is the accuracy difference of PRS between using multi-ancestry GWAS ($PRS_{multi}$) and using BBJ GWAS ($PRS_{Minor\_GWAS}$) when the number of bins from EUR GWAS is 64. The x-axis is the number of bins from BBJ included in the multi-ancestry GWAS. The error bars indicate the standard error of mean accuracy improvement. The red dashed line is y=0. The traits were ranked by polygenicity estimates using UKBB (**Figure 3**).

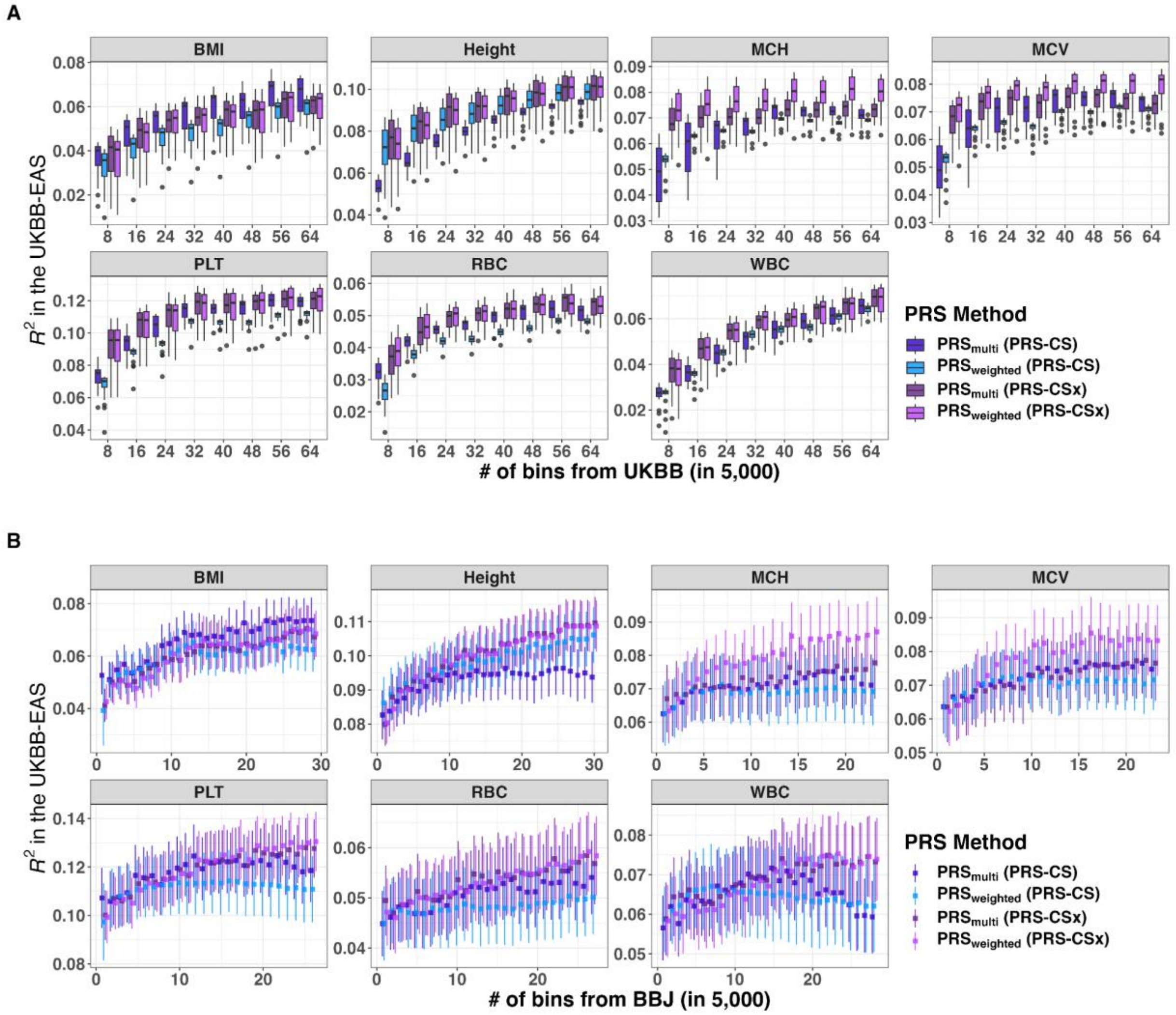

#### Figure S12. Predictive accuracy using PRS-CS and PRS-CSx as a function of sample size in the UKBB-EAS.

We constructed PRS using PRS-CS and PRS-CSx, and evaluated them in the UKBB-EAS. We explored the impact of sample sizes from EUR GWAS (A) and Minor GWAS (B) on PRS accuracies. While in B), we illustrated the results using 64 bins from EUR GWAS as an example. The error bars are the standard errors of predictive accuracy. We showed the results for 7 traits with SNP-based heritability > 0.1 in both BBJ and UKBB, while they were ranked by polygenicity estimates using UKBB (**Figure 3**). Full results are shown in **Table S8** and **Table S9**.

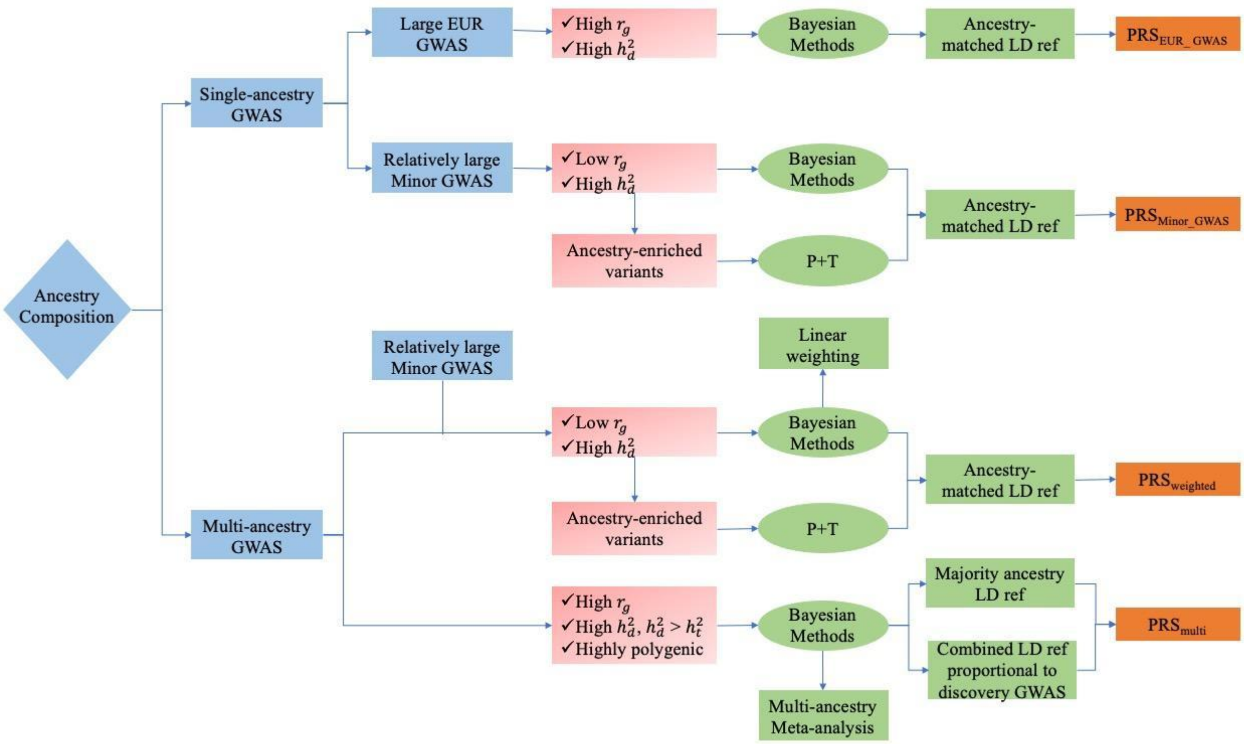

#### Figure S13. Flow chart for best practices using single-ancestry and multi-ancestry GWAS in PRS analyses.

We summarized the key considerations in the development of PRS. The factors taken into account include the ancestry composition of discovery GWAS (blue), the trait-specific genetic architecture (pink) and PRS methodology (green). The development of single-ancestry PRS involves $PRS_{EUR\_GWAS}$ derived from European GWAS (EUR GWAS), and $PRS_{Minor\_GWAS}$ derived from GWAS conducted on minority populations (Minor GWAS). Multi-ancestry PRS include $PRS_{multi}$, which is derived from meta-analyzed multi-ancestry GWAS, and $PRS_{weighted}$, obtained from a linear combination of ancestry-specific PRS. Bayesian methods, such as PRS-CS and PRS-CSx, adapt to trait-specific genetic architecture, are recommended. Alternative methods that jointly estimate effect sizes while accounting for linkage disequilibrium can also be considered. Abbreviations: Cross-ancestry genetic correlation ($r_g$), SNP-based heritability in discovery ($h_d^2$) and target populations ($h_t^2$).

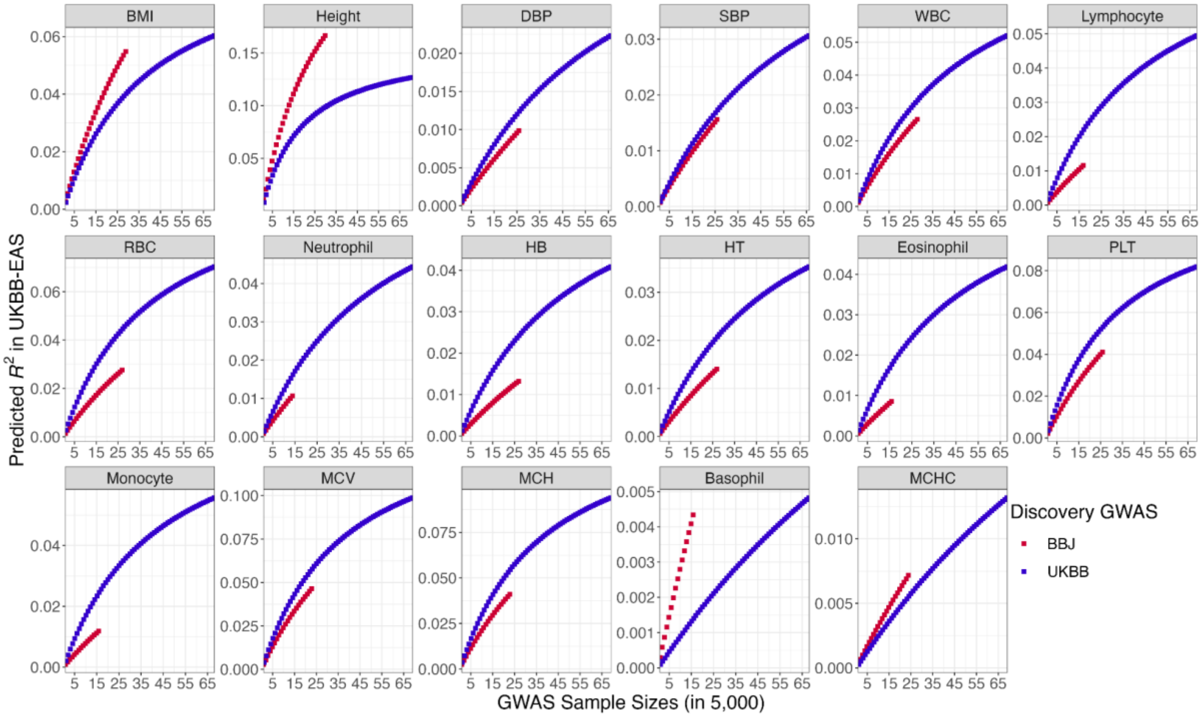

#### Figure S14. Predicted accuracy in the UKBB-EAS using different discovery GWAS.

The predicted accuracy of PRS in the UKBB-EAS using GWAS from BBJ and UKBB, respectively, was calculated using the theoretical equation described in detail in **Supplementary Note 8**.

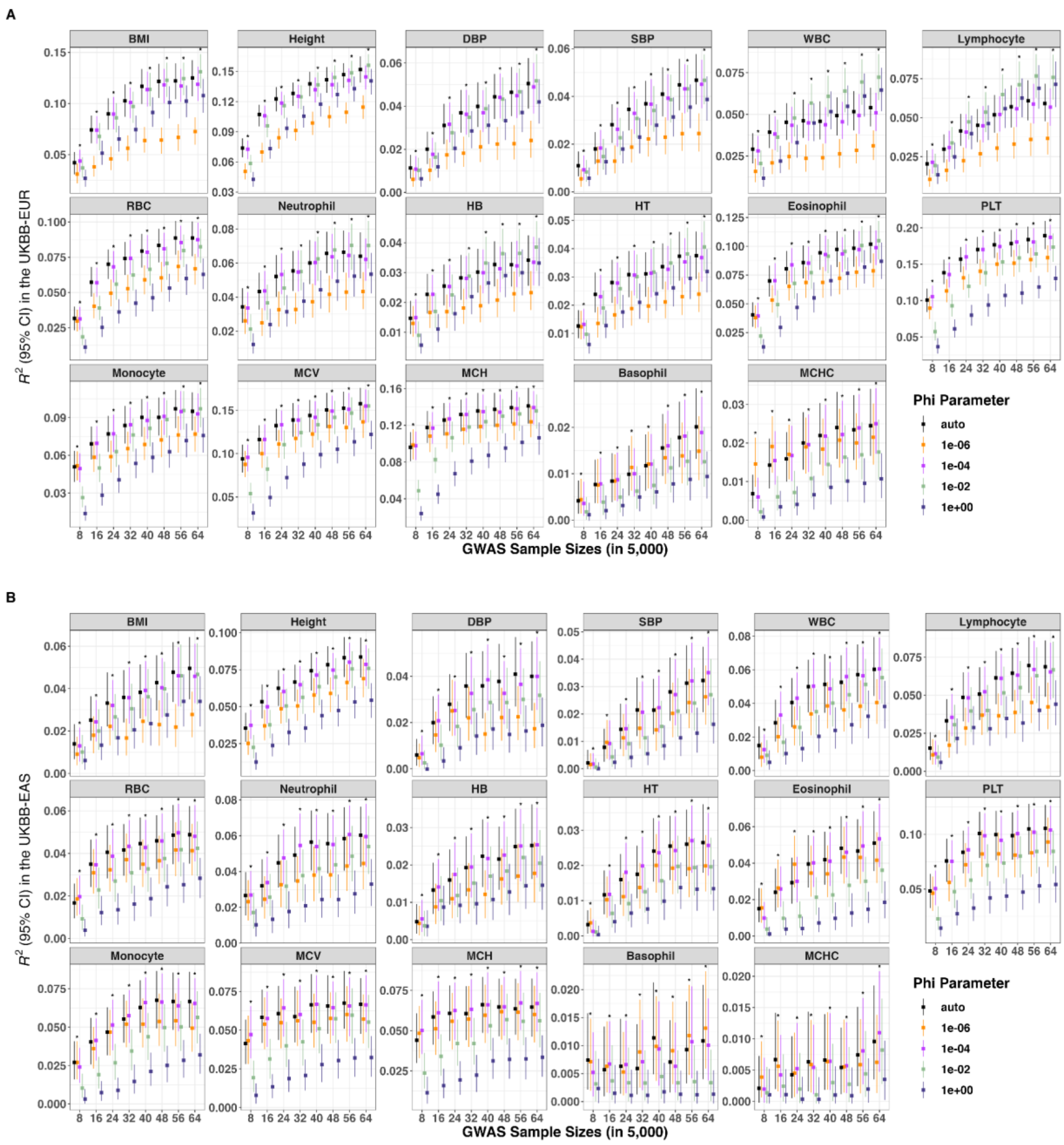

### Figure S15. Prediction performance of PRS-CS using different models in the UKBB.

We varied the phi parameters for PRS-CS to construct PRS in the A) UKBB-EUR and B) UKBB-EAS. The asterisks indicated the optimal phi parameters achieving highest predictive accuracy in the grid model. We found that the auto model which directly estimated the phi parameter from input GWAS performed comparable to the grid model which required fine-tuning.

#### Supplementary Tables

Table S3: Descriptions of 17 studied phenotypes in the UK Biobank (UKBB) and Biobank Japan (BBJ).

| Phenotypes | Abb. | UKBB phenocode | No. of total Bins in the UKBB | No. of total Bins in the BBJ |
|---|---|---|---|---|
| basophil count | Basophil | 30160 | 68 | 16 |
| body mass index | BMI | 21001 | 70 | 29 |
| diastolic blood pressure | DBP | 4079 | 66 | 26 |
| eosinophil count | Eosinophil | 30150 | 68 | 16 |
| height | Height | 50 | 71 | 30 |
| hematocrit percentage | HT | 30030 | 69 | 27 |
| hemoglobin concentration | HB | 30020 | 69 | 27 |
| lymphocyte count | Lymphocyte | 30120 | 68 | 17 |
| mean corpuscular hemoglobin | MCH | 30050 | 69 | 23 |
| mean corpuscular hemoglobin concentration | MCHC | 30060 | 69 | 24 |
| mean corpuscular volume | MCV | 30040 | 69 | 23 |
| monocyte count | Monocyte | 30130 | 68 | 16 |
| neutrophil count | Neutrophil | 30140 | 68 | 14 |
| platelet count | PLT | 30080 | 69 | 26 |
| red blood cell count | RBC | 30010 | 69 | 27 |
| systolic blood pressure | SBP | 4080 | 66 | 26 |
| white blood cell count | WBC | 30000 | 69 | 28 |

Table S4. Genetic architecture of 17 studied phenotypes.

| Phenotype | Polygenicity (SD) | | SNP-based heritability (SD) | | *S* (SD) | |
|---|---|---|---|---|---|---|
| | **UKBB** | **BBJ** | **UKBB** | **BBJ** | **UKBB** | **BBJ** |
| BMI | 0.04778 (0.00165) | 0.02256 (0.00142) | 0.259 (0.002) | 0.167 (0.003) | -0.593 (0.028) | -0.39 (0.067) |
| Basophil | 0.00369 (0.00027) | 0.00065 (0.00018) | 0.049 (0.001) | 0.054 (0.005) | -0.718 (0.057) | -0.594 (0.151) |
| DBP | 0.02191 (0.00095) | 0.00647 (0.00124) | 0.14 (0.002) | 0.067 (0.003) | -0.533 (0.04) | -0.643 (0.098) |
| Eosinophil | 0.00703 (0.00022) | 0.00202 (3e-04) | 0.203 (0.002) | 0.077 (0.005) | -0.634 (0.036) | -0.38 (0.14) |
| HB | 0.01062 (0.00033) | 0.00666 (0.00062) | 0.184 (0.002) | 0.077 (0.003) | -0.632 (0.035) | -0.544 (0.092) |
| HT | 0.01036 (0.00033) | 0.00599 (0.00054) | 0.175 (0.002) | 0.079 (0.003) | -0.626 (0.035) | -0.49 (0.09) |
| Height | 0.02256 (0.00042) | 0.01246 (0.00046) | 0.539 (0.002) | 0.333 (0.004) | -0.687 (0.02) | -0.427 (0.043) |
| Lymphocyte | 0.0119 (0.00036) | 0.00659 (0.00906) | 0.207 (0.002) | 0.088 (0.004) | -0.581 (0.034) | -0.73 (0.15) |
| MCHC | 0.00206 (0.00016) | 0.00079 (0.00013) | 0.053 (0.001) | 0.058 (0.003) | -0.665 (0.066) | -0.511 (0.116) |
| MCH | 0.00441 (0.00015) | 0.00125 (1e-04) | 0.256 (0.002) | 0.156 (0.003) | -0.615 (0.04) | -0.253 (0.093) |
| MCV | 0.00525 (0.00017) | 0.00185 (0.00014) | 0.264 (0.002) | 0.167 (0.003) | -0.633 (0.038) | -0.36 (0.081) |
| Monocyte | 0.00601 (0.00019) | 0.00242 (0.00032) | 0.23 (0.002) | 0.092 (0.005) | -0.453 (0.039) | -0.412 (0.121) |
| Neutrophil | 0.01077 (0.00036) | 0.00287 (0.00041) | 0.179 (0.002) | 0.093 (0.005) | -0.689 (0.032) | -0.457 (0.129) |
| PLT | 0.00682 (0.00019) | 0.00405 (0.00027) | 0.299 (0.002) | 0.148 (0.003) | -0.581 (0.034) | -0.457 (0.065) |

| | | | | | | |
|---|---|---|---|---|---|---|
| RBC | 0.01081 (0.00031) | 0.00363 (0.00028) | 0.247 (0.002) | 0.116 (0.003) | -0.615 (0.03) | -0.44 (0.076) |
| SBP | 0.02131 (0.00086) | 0.00928 (0.00089) | 0.151 (0.002) | 0.086 (0.003) | -0.568 (0.038) | -0.409 (0.108) |
| WBC | 0.01414 (0.00042) | 0.00696 (5e-04) | 0.205 (0.002) | 0.111 (0.003) | -0.639 (0.03) | -0.518 (0.076) |
